# Range geography and temperature variability explain cross-continental convergence in range and phenology shifts in a model insect taxon

**DOI:** 10.1101/2024.07.22.604548

**Authors:** Catherine Sirois-Delisle, Susan C.C. Gordon, Jeremy T. Kerr

## Abstract

Climate change may introduce conditions beyond species’ tolerances; to survive, species must avoid these extremes. Phenological shifts are one strategy, as species move their activity or life history events in time to avoid extreme conditions. Species may also shift in space, moving their ranges poleward to escape extremes. However, whether species are more likely to exhibit one or both strategies, and whether this can be predicted based on a species’ functional traits, is unknown. Using a powerful macroecological dataset of European and North American odonate observations, we assessed range and phenology shifts between two time periods (1980-2002 and 2008-2018) to measure the strength and direction of the association between responses. Species with the greatest poleward range shifts also showed the largest phenological shifts toward earlier annual activity periods, with half of all species shifting in both space and time. This response was consistent across continents, despite highly divergent land use and biogeographical histories in these regions. Surprisingly, species’ range and phenology shifts were not related to functional traits; rather, southern species shifted their range limits more strongly, while increasing temperature variability hindered range shifts. By reducing risk through phenological shifts, the resulting larger populations may be more likely to disperse and expand species’ ranges. Species shifting in both space and time may be more resilient to extreme conditions, although further work integrating abundance data is needed. We also identified a small number of species (approximately 10%) that failed to shift at all; these species are likely to be particularly vulnerable to climate change, and should be prioritized for conservation intervention.

## Introduction

Climate change alters climatic means and increases the frequency of extreme weather events, exposing species to conditions outside of their tolerances, and often leading to population declines (Goulson, 2019; IPCC, 2021). Species may avoid extreme conditions by dispersing to new areas where conditions pose fewer weather-related challenges, often leading to poleward range expansion (Davis et al., 2005; Lawlor et al., 2024). Species’ biological timing could also shift, through adaptation or phenotypic plasticity, with earlier warming advancing the timing of early-season activities and life-history events (Davis et al., 2005; Hällfors et al., 2024; Novella-Fernandez et al., 2023; Parmesan and Yohe, 2003). Both species’ geographical ranges and seasonal timing depend strongly on climate and habitat conditions, with shifts in space and time permitting species to remain within the limits of their ecological niches (Chen et al., 2011; Engelhardt et al., 2022; Grewe et al., 2013; Menzel et al., 2006; Parmesan and Yohe, 2003). This allows populations to grow, despite changing environments, and reduces the risk of climate debt and extinction (Devictor et al., 2012; Franks et al., 2018; Lustenhouwer et al., 2018; Saino et al., 2010; Souza et al., 2023; Urban, 2015).

Positive population trends can be stronger in species that shift both their range and phenology (Hällfors et al., 2024). Greater phenological plasticity under warmer spring temperatures may increase reproductive success, leading to greater population growth and range expansions (Macgregor et al., 2019), as positive or stable trends in species abundance and habitat availability are essential for range shifts (Mair et al., 2014; Platts et al., 2019). However, there could also be a trade-off between phenological and geographical shifts (Amano et al., 2014; Hassall, 2015; Socolar et al., 2017). Species with greater dispersal abilities may have less need for phenological shifts as they track their climatic niche through space, while weaker dispersers may be confronted with greater selective pressure to shift phenology within their range (Amano et al., 2014; Hassall, 2015; Socolar et al., 2017). Since range shifts can also result from extirpations at species’ trailing range edge (Parmesan et al., 1999), greater phenological shifts may mitigate the need for range shifts, as species better tolerate new climatic conditions. Cross continental studies report converging effects of climate change on species’ range shifts and abundances (Neate-Clegg et al., 2024; Stephens et al., 2016), including among insects (Kerr et al., 2015; Neate-Clegg et al., 2024; Pinkert et al., 2022), but potential relationships between phenological and geographic responses have not yet been investigated at continental scales. Functional traits, such as dispersal, may determine species’ spatial and temporal responses to climate change (Chen et al., 2011; Kharouba et al., 2009; Schuetz et al., 2019; Zografou et al., 2021). However, these relationships are inconsistent across taxa and regions, and cross-continental tests have not been attempted (Angert et al., 2011; Buckley and Kingsolver, 2012; Estrada et al., 2016; MacLean and Beissinger, 2017). Geographic locations and environmental characteristics of species’ ranges may also predict range shifts, as animal species with high latitude ranges have been shown to exhibit smaller range shifts (MacLean and Beissinger, 2017; Pinkert et al., 2022), while increasing local temperature and loss of natural landcover may drive range retractions (Pacifici et al., 2020). However, exposure to extreme climate events (e.g. drought, heat waves, or storms) within species ranges may disrupt species’ dispersal abilities and capacities to tolerate new conditions (Kerr, 2020; Román-Palacios and Wiens, 2020). Exposure to thermal anomalies can rapidly change entire communities and create shifts towards new ecosystems, sometimes leading to local declines (Day et al., 2018; Grant et al., 2017; Harris et al., 2018; Román-Palacios and Wiens, 2020).

While global change research on insects often emphasizes butterfly and bee taxa, recently assembled databases of odonate observations provide a rare opportunity to investigate species’ spatiotemporal responses at larger taxonomic and spatial scales, particularly as most work has been done at national scales (Córdoba-Aguilar et al., 2023; Kalkman et al., 2018; Sandall et al., 2022). Due to their use of aquatic and terrestrial habitat across life different stages, dragonflies and damselflies are also considered indicator species for both terrestrial and aquatic insect responses to changing climates (Hassall, 2015; Pinkert et al., 2022; Šigutová et al., 2025), giving the study of these species broad relevance for conservation. There is some evidence that functional traits relate to odonates’ interspecific variation in range shifts (Angert et al., 2011; Grewe et al., 2013), phenology shifts (Diamond et al., 2011; Gutiérrez and Wilson, 2021; Zografou et al., 2021), extinction risks (Cardillo et al., 2008; Cooper et al., 2008; Rocha-Ortega et al., 2022; Suhonen et al., 2022), and rates of decline and expansion within limited geographic scopes (Powney et al., 2015; Rapacciuolo et al., 2017; Rocha-Ortega et al., 2020). While relationships between morphological traits and range boundaries have been shown for some groups (i.e. Rundle et al., 2007), these may depend on species’ geographic context. For example, differences in habitat connectivity and dispersal ability may constrain range shifts for lentic species (those species that breed in slow moving water like lakes or ponds) and lotic species (those living in fast moving-water) in different ways (Kalkman et al., 2018). More southerly lentic species may expand their range boundaries more than lotic species, as species accustomed to ephemeral lentic habitats better dispersers (Grewe et al., 2013), yet lotic species have also been found to expand their ranges more often than lentic species, potentially due to the loss of lentic habitat in some areas (Bowler et al., 2021). While warm-adapted species with more equatorial distributions could expand their ranges poleward following warming (Devictor et al., 2008), they could also increase in abundance in this new range area relative to species that historically occupied those areas and are less heat-tolerant (Powney et al., 2015).

In this study, we tested whether species with stronger geographical range shifts also advanced their emergence phenologies, or if one response offsets the need for the other. We also asked whether functional traits, range geography (i.e. southerly vs. northerly), or temperature variability predict range shifts at species’ northern range limits, and whether these factors can also predict shifts in species emergence phenology. We predicted that species would exhibit shifts in both geography and phenology, as we expect species shifting in phenology to have larger populations, increasing likelihood of dispersal and successful range shifts. We predicted that species able to use both lentic and lotic habitats would shift their phenologies and geographies more than those able to use just one habitat type, as generalists outperform specialists as climate and land uses change (Ball-Damerow et al., 2015, 2014; Hassall and Thompson, 2008; Powney et al., 2015; Rapacciuolo et al., 2017). Alternatively, species might respond to rapid climate changes in ways that reflect their geographical position, indicating that where they are found is a better predictor of their conservation risk from global change than their intrinsic biological characteristics.

**Figure 1:**
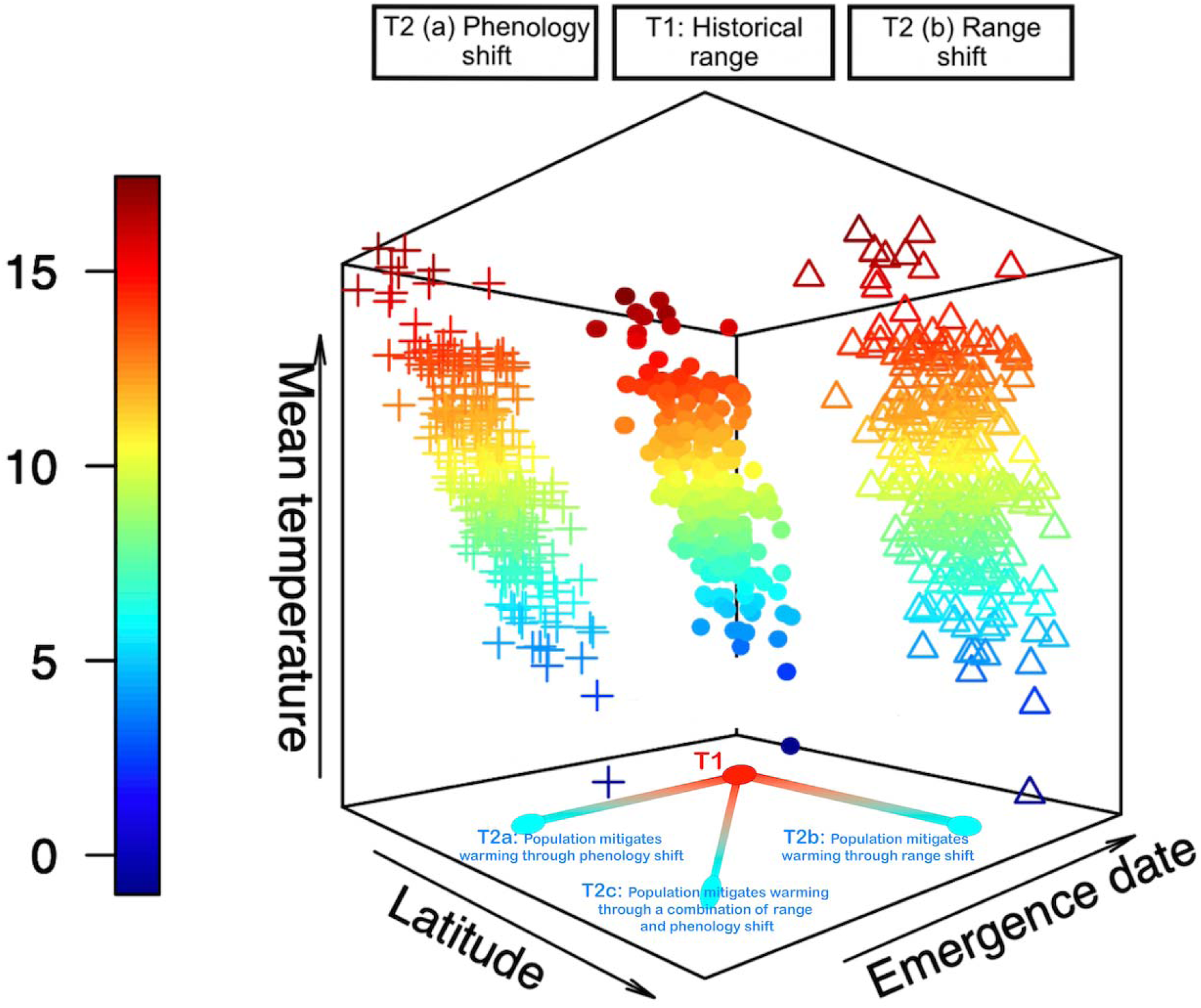
Representation of temporal and geographical limits characterising the ecological niche of a hypothetical odonate species. Points show 250 individuals according to their Julian day of emergence, latitudinal position, and temperatures to which each individual is exposed. Points represent historical observations (T1), plus signs show observations following a shift towards earlier emergence dates after warming. (T2a) triangle symbols show observations following a shift towards higher latitudes after warming (T2b). Species could also shift both range and phenology in response to warming (T2c). Warm and cool colors show hot and cold temperatures, respectively.

## Methods

### Biological records

We assembled ∼2 million observations records for North American and European odonate species collected between 1980 and 2018. Data sources included online dataset aggregators GBIF (http://gbif.org/) and Canadensys (http://www.canadensys.net/), Odonata Central (Abbott, 2020), and other institutions (see Acknowledgments). While odonates were sampled opportunistically, biases associated with data that are not systematically collected are less likely to affect trends at large spatial and temporal scales, particularly if data are obtained from multiple independent sources (Pyke and Ehrlich, 2010; Zattara and Aizen, 2021). We removed records with incomplete or missing species identification, year, or locality information. We selected unique observations for species, location, year, and Julian day of collection, and restricted the data to continental North America and Europe. We mapped species-specific observations using ArcGIS software (ESRI, 2019), and qualitatively verified species ranges. If a species was found on both continents, we only retained observations from the continent that was the most densely sampled. If we merged data for one species found on both continents, we could not perform a cross-continental comparison. However, if the same species on different continents was treated as different species, this would lead to uninterpretable outcomes (and the creation of pseudo-replication) in the context of phylogenetic analyses. In addition, species found on both continents did not have sufficient data to meet criteria for the phenology analysis.

We followed widely accepted methods to determine species range boundaries (Devictor et al., 2012, 2008; Kerr et al., 2015), although other methods exist that are appropriate for different data types and research questions i.e. (Ni and Vellend, 2021). We assigned species presences to 100×100 km quadrats, a scale that is large enough to maintain adequate sampling intensity but still relevant to conservation and policy (Soroye et al., 2020), to identify the best sampled species. We excluded species found in fewer than 50 quadrats to increase the likelihood of accurately predicting the position of species’ northern range boundaries. We retained ∼1,100,000 records from Canada, the United States, and Northern Mexico, comprising 76 species (Figure 2). Observation records were separated into two time periods, to compare species’ recent phenologies and northern range limits (2008 to 2018) to conditions in a historical time period (1980 to 2002).

**Figure 2:**
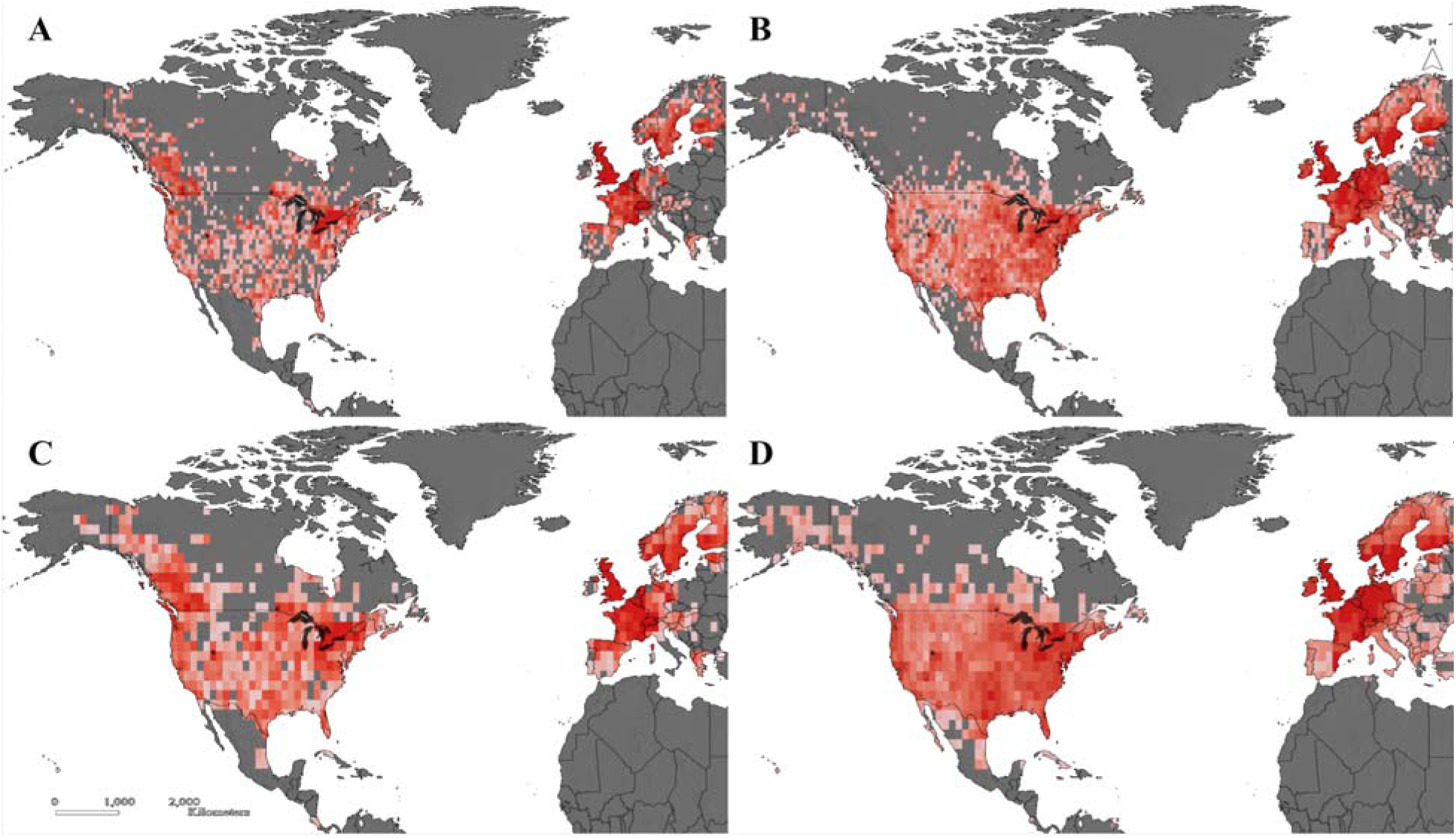
Richness of 76 odonate species sampled in North America and Europe in the historic period (1980-2002; panes A and C) and the recent period (2008-2018; panes B and D). Species richness per 100 × 100 km quadrat is shown in panes A and B, while panes C and D show species richness per 200 × 200 km quadrat. Dark red indicates high species richness, while light pink indicates low species richness.

### Temperature Variability

Temperature variability during a species’ flight season may impact its ability to establish in new locations, or shift its emergence timing the following year. We downloaded a high-resolution gridded dataset for monthly average daily maximum temperature from the Climatic Research Unit (New et al., 2002). We extracted average values per 100×100 km quadrat across the months of April to October, covering the main flight period for odonates in this study, for each year included in the historical and recent periods. We calculated the coefficient of variation per quadrat for each time period and averaged these values per species and time period to measure interannual temperature variability during species’ flight season across their range. We used the difference between recent and historical measures as an estimate of change in temperature variability.

### Spatial and temporal change metrics

To limit potential effects of temporal and spatial biases, we generated range and phenology shift metrics with specific criteria for quadrats and species selection (Bartomeus et al., 2018; Gaiji et al., 2013; García-Roselló et al., 2015); we retained 76 species for range shift estimates (Supplementary Information). Species’ northern range boundaries were calculated using the mean of the 10 most northern points of each species range in both the historic (1980 – 2002) and recent (2008 – 2018) time periods, measured in kilometers from the equator (as in Kerr et al., 2015). We used the difference between range limit positions in the historic recent time periods to estimate species’ northern range limit shifts.

Since spatiotemporal biases sometimes inflate range shift measurements (Kujala et al., 2013), we used null models to test whether observed range shift estimates were robust, and the extent to which those responses differed from expectations arising because of rising sampling intensity over time. 1000 randomized datasets were created with the same number of species-specific geographical points per time period as the number of actual observations. Maximum and minimum latitude and longitude values were held constant relative to observed values. Northern range limit shifts were calculated as the difference in the mean of the 10 most northern points between the historic and recent time periods. We used generalized linear models (GLMs; glm command in R) to test whether species-specific range limit shifts in each iteration predicted range shifts measured from the observation data (Figure S1, S2). We found that observed northern range limit shifts are not consistent with expectations derived from changes in sampling intensity.

We estimated species-specific emergence phenology for each time period in 200×200 km quadrats; using larger quadrats increases probabilities of detecting signals of emergence phenology, which may otherwise be lost due to gaps in data density. We retained quadrats that contained at least 25 observations for a given species in both time periods. To estimate phenology per area, we used the *weib.limit* function of the *WeibullR* R package (Pearse et al., 2017). This function uses the Weibull distribution to estimate the Julian day of a species’ first appearance, and is especially useful to measure the timing of phenological events in sparsely sampled datasets. Techniques such as using the average of the n^th^ first observations of phenological events, or the n^th^ percentile flight dates (Brooks et al., 2014; Robbirt et al., 2011), tend to overestimate the timing of biological events due to temporal bias towards later days in the species’ active period (Pearse et al., 2017). We retained emergence estimates between March 1^st^ and September 1^st^, as well as species and quadrats that showed a difference in emergence phenology of −25 to 25 days, −30 to 30 days, or −35 to 35 days between both time periods, to include only phenology shifts that could be biologically meaningful to environmental climate change. 68 species found across 63 quadrats met these criteria. Large changes in phenology are likely explained by other anthropogenic or natural factors, or could occur due to noise in the data, since these phenology calculations per region are extremely data intensive. We calculated the difference in the day of emergence per quadrat between both time periods, as well as mean phenology change across all quadrats for each species. The number of quadrats per species used to calculate their mean phenological shift varied between 2 and 46.

We used null models to assess whether our approach to estimating phenology shifts was robust, as potential issues may arise due to spatiotemporal biases in the underlying data (Kujala et al., 2013). We constructed 1000 randomized datasets of species’ hypothetical days of occurrence, using the same number of quadrats, and observations within quadrats, as in each time period of the observation records. We assigned the maximum and minimum Julian day of occurrence from the observation records to limit values in the randomized datasets. We applied the same method and criteria of inclusion to the randomized datasets as we did to measure phenology shifts from the observation data. GLMs were built to test whether phenology shifts calculated using 1000 random datasets predicted the phenology shifts that we measured. No discernable pattern emerged, indicating that observed shifts in phenology are not consistent with expectations derived from differences in sampling intensity over time (Figure S3, S4).

### Range geographies and functional traits

To assemble trait data for the 76 species in the database, we used field guides (Cannings, 2002; Jones et al., 2008; Paulson, 2012) and existing trait databases (Powney et al., 2014; Waller et al., 2019). We considered any evolved morphological, physiological, behavioural, or life history characteristic as a functional trait (Beissinger and Riddell, 2021). Geographic range and associated climatic characteristics are often considered ecological traits, as they are consequences of functional traits and their interactions with geographic features (Bried and Rocha-Ortega, 2023; Chichorro et al., 2019). Such ecological variables may predict species’ responses to climate change, and can add significant value to predictive models (MacLean and Beissinger, 2017). We identified whether species’ ranges were more northern, southern, or both northern and southern (both), and determined the range size of each species by counting the number of quadrats occupied by that species in the historical time period.

Along with the geographic and climatic attributes, temperature variability and distribution, we selected four functional traits likely to be biologically relevant to spatial and temporal responses to climate change: flight duration, breeding habitat type (lotic, lentic, or both), egg laying habitat (exophytic vs. endophytic), and body size (Cannings, 2002; Jones et al., 2008; Paulson 2012; Powney et al., 2014; Waller et al., 2019). Species’ flight period was measured as the total number of days of the flight period, estimated from the approximate time of the month of average first and last appearances. Breeding habitat was assigned according to a species’ uses of lotic, lentic, or both habitat types. Egg laying habitat was assigned according to whether species use exophytic egg-laying habitat (i.e. eggs laid in water or on land, relatively larger in number), or endophytic egg-laying habitat (i.e. eggs laid inside plants, usually fewer in number); species using exophytic habitats are associated with greater northward range limit shifts (Angert et al., 2011). Body size corresponded to the mean length of the abdomen of each species. We excluded overwintering stage and range size from our analysis as data were incomplete for many species, and excluded migratory behavior as the vast majority of species included in the study were non-migratory.

We tested for correlations amongst all predictors by calculating the Predictive Power Statistic (PPS) and Pearson correlations among traits (van der Laken, 2021): we found no evidence of correlation.

### Statistical analyses

We conducted statistical analyses using R Statistical Software (R Core Team, 2019). All continuous variables were transformed into Z-scores using the *scale* function in R. First, we investigated whether there was a relationship between species’ range and phenological shifts by modelling phenology shift as the dependent variable, and range shift and continent as independent variables. We used both species-level frequentist (GLM; glm function in R) and Bayesian (Markov Chain Monte Carlo generalized linear mixed model, MCMCglmm; Hadfield, 2010) models to improve the robustness of the results. We included a term to account for phylogeny in the MCMCglmm model, as species that are closely related are likely to have similar traits. We used the molecular phylogenetic tree published by the Odonate Phenotypic Database (Waller et al., 2019), which used a morphological and taxonomic phylogeny as the backbone tree, allowing species to move within their named genera or families according to molecular evidence (Waller and Svensson, 2017). Trace and density plots for the MCMCglmm model revealed no issues with autocorrelation or model convergence (Figure S7).

Next, we investigated whether functional traits, range geography, or temperature variability predicted range shifts at species’ northern range limits, and whether the same predictors explaining range expansions could also predict shifts in species emergence phenology. We constructed two sets of GLMs, in addition to two sets of MCMCglmms accounting for phylogeny; one of each with changes in species’ northern range limits as the response variable, and the other with changes in emergence phenology as the response variable. Non-significant variables, specifically all functional traits, were removed from the final geographic range shift model. No effects were significant in the model of phenology shifts. Trace and density plots for the MCMCglmm models did not indicate limitations related to autocorrelation or model convergence (Figure S8).

In addition to the inclusion of phylogeny in statistical models to account for potential data non-independence, we measured the phylogenetic signal in range and phenological shifts. We used the *phylosig* function of the *Phytools* package version 0.7-70 (Revell, 2012), which calculates phylogenetic signal using Pagel’s lambda and Blomberg’s K.

## Results

### Relationship between range and phenology shifts

Most species (52 of 76) expanded their northern range limits toward higher latitudes (mean range expansions of 180 km). The average range expansion across all species was 63 km northward, although some species showed range retractions (Figure 4). Most species (41 of 66) maintained or advanced their emergence phenology (mean of −2.71 days in emergence phenology shifts among all species; Figure 4). Fewer species were included in phenology analyses than analyses of range shifts due to the data intensity required to capture phenology shifts at a study site (N=66 vs. N = 76). Many species (50%) showed both advancing emergence phenology and range expansions, while 10% of species showed neither range nor phenological shifts relative to historical baselines.

The effect of species’ range shifts on phenology range shifts was significant in our model investigating the relationship between these responses, indicating that species shifting their northern range limits to higher latitudes also showed stronger advances in their emergence phenology (Figure 3). This result that was consistent in GLM and Bayesian analyses (p < 0.01; Table 1), and was maintained across North America and Europe, with no effect of continent in the model. This trend was consistent among both dragonflies and damselflies, although there was considerable interspecific variation in the magnitude of spatial and temporal shifts. Accounting for phylogeny did not improve model predictions and did not explain greater model variance (R^2^ = 17% and 15%, for GLM and MCMCglmm models, respectively).

**Figure 3:**
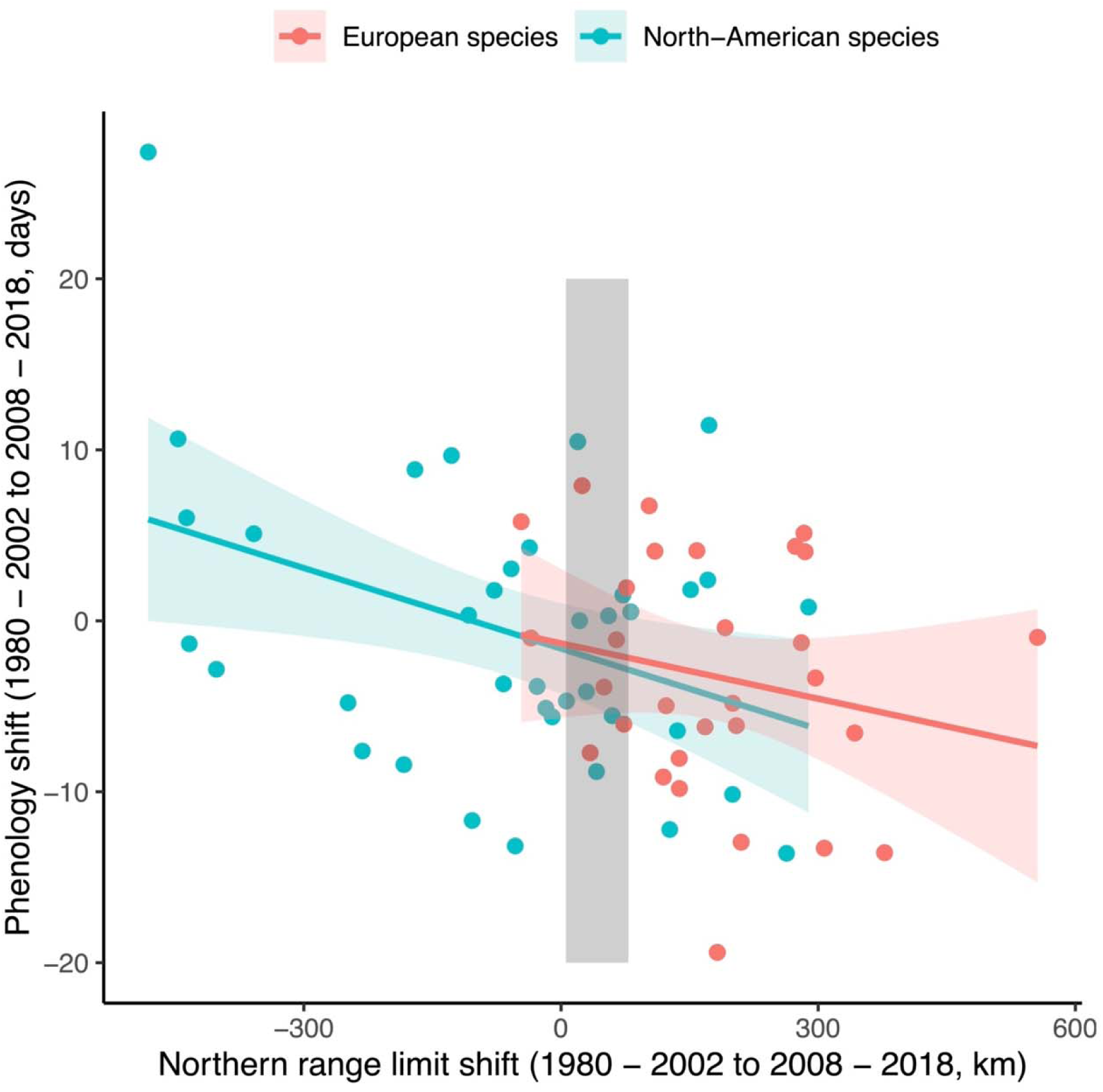
Relationship between range shifts and emergence phenology shifts among North American and European odonate species (N = 66; model R^2^ = 17.08 for glm, 14.9% for MCMCglmm). For reference, the shaded area shows mean latitudinal range shifts of terrestrial taxa as reported by Lenoir et al. (calculated as the yearly mean dispersal rate of 1.11 +/− 0.96 km per year over 38 years).

**Table 1:** Fixed effects estimates and associated statistics from the generalized linear model and generalized mixed effects model (accounting for phylogeny; for credible intervals, see Table S4) of the relationship between range shifts and emergence phenology change. The continent term shows effects of the North American continent compared to the European continent as the reference level. N gives the number of species involved in the model, and an asterisk indicates statistical significance of the variable in question (p-value < 0.05). The pseudo R^2^ type is Nagelkerke (Nagelkerke, 1991).

| Predictors | Phenology shift<br>(N = 66) |  |  |  |
| --- | --- | --- | --- | --- |
|  | GLM |  | MCMCglmm |  |
|  | Est. | p | Post.m | p |
| (intercept) | 0.12 | 0.53 | 0.12 | -0.49 |
| Range shift | -0.45 | <0.01* | -0.45 | <0.01* |
| Continent | -0.22 | 0.44 | -0.22 | 0.42 |
| **Model evaluation** |  |  |  |  |
| AIC/DIC | 185.39 |  | 185.43 |  |
| Null model | 193.13 |  | 193.12 |  |
| Pseudo R<sup>2</sup> | 17.08% |  | 14.90% |  |

### Drivers of range and phenology shifts

Range geography and climate variability, rather than functional traits, predicted range shifts in both North America and Europe, with range geography being consistently the strongest predictor. Species’ functional traits did not relate to the extent of observed geographical range shifts in tests using GLM and MCMCglmm models. Species with more southern distributions shifted their northern range limits towards higher latitudes more than northern species or species present in both the north and south (Table 2; p = 0.002 and 0.004, model R^2^ = 26.6% and 23.7%, for GLM and MCMCglmm models, respectively), with no effect of range size on range shifts. Species experiencing smaller changes in interannual temperature variability also had a higher likelihood of northern range limit shifts (p = 0.0005 for GLM, p = 0.002 for MCMCglmm). Results from the GLM and MCMCglmm models were qualitatively similar, however, a smaller amount of model variance was explained when phylogeny was accounted for. Emergence phenology shifts were not affected by species’ traits, range geography, nor climate variability; due to this, model results are not displayed here. While range and phenology response types are related, this suggests that the mechanisms underlying phenological shifts are different than those underlying range shifts.

**Table 2:** Fixed effects estimates and associated statistics from the generalized linear model and generalized mixed effects model (accounting for phylogeny; for credible intervals, see Table S4) of drivers of odonate range shifts. N indicates the number of modelled species, an asterisk indicates statistical significance of the variable in question, and a dash symbol shows that the variable was excluded from the final model. The pseudo R^2^ type is Nagelkerke (Nagelkerke, 1991). For the categorical variables breeding habitat type and range geography, we used lotic habitat type and Northern range as reference levels, respectively.

|  | Range shift (N = 76) |  |  |  |
| --- | --- | --- | --- | --- |
|  | GLM |  | MCMCglmm |  |
| Predictors | Est. | p | Post.m | p |
| (intercept) | -0.65 | 0.018 | -0.65 | 0.022 |
| Widespread distribution | 0.34 | 0.32 | 0.34 | 0.31 |
| Southern distribution | 0.95 | 0.002 | 0.95 | 0.004 |
| T ° variability | -0.38 | 0.0005 | -0.38 | 0.002 |
| Model evaluation |  |  |  |  |
| AIC/DIC | 202.8 |  | 202.9 |  |
| Null model | 218.7 |  | 218.7 |  |
| Pseudo R<sup>2</sup> | 26.60% |  | 23.70% |  |
| Phylogenetic signal |  |  |  |  |
| Pagel's lambda (p) | 0.0057 (0.89) |  |  |  |
| Blomberg's K (p) | 0.11 (0.47) |  |  |  |

A phylogenetic signal may indicate that there are traits that determine species’ spatial and temporal responses to changing climate that were not measured in this study. Yet we detected no phylogenetic signal using Pagel’s lambda or Blomberg’s K in either geographical range or phenological responses (Table 2). Adding a phylogeny term to the MCMCglmm models also failed to produce a pattern different to the GLMs, and model performance did not improve when we accounted for phylogeny in our assessment of northern range shifts.

## Discussion

In one of the first studies to investigate both shifts of phenology and range at a continental scale, we find that dragonfly and damselfly species show pronounced geographical and phenological shifts that converged across Europe and North America. Species expanding their ranges poleward also emerged earlier in the spring on both continents (Figure 3), with shifts predicted by range geography and climate variability, but not functional traits. These results suggest that some species may have an advantage with respect to climate change: they demonstrate the flexibility to respond both temporally and spatially to the onset of rapid climate change. Conversely, species that show neither geographic nor phenological shifts may be particularly vulnerable to climate change.

We found no evidence for a tradeoff between range and phenology shifts; instead, half of species shifted both range and phenology. Earlier seasonal timing allows species to stay within their climatic limits and maintain population growth rates (Macgregor et al., 2019), although earlier emergence could expose individuals to early season weather extremes (McCauley et al., 2018). As only a small proportion of odonate adults undertake long range dispersal (Conrad et al., 1999), greater local population sizes should contribute to higher dispersal rates (Mair et al., 2014), facilitating range shifts (Kerr, 2020; Leroux et al., 2013). This is consistent with results from other taxa: among British butterflies, early emergence increased population growth and facilitated range shifts for species with multiple generations per year (Macgregor et al., 2019); Finnish butterfly species with the greatest population growth rates shifted both their phenology and ranges (Hällfors et al., 2021). Such population growth or maintenance, and therefore the potential for range shifts, is only possible if habitat is available (Mair et al., 2014). Future work should consider habitat availability alongside range and phenology shifts, as it may help explain why some species are able to shift their phenology but not their range.

Southern species were more likely to expand their ranges northward than northern species or species present in both the north and south. Species’ ability to maintain large populations may be impaired in northern latitudes, where rates of climate change are high (IPCC, 2021), hindering dispersal and colonization that are precursors to range expansions (Mair et al., 2014). Further mechanistic understanding of these processes requires abundance data. Southern species may have narrower niche breadths than widespread or northern species, and may respond more rapidly to climate change to track this narrower niche (Hällfors et al., 2024). Emerging mean conditions in areas adjacent to the ranges of southern species may offer opportunities for range expansions of these relative climate specialists, which can then tolerate climate warming in areas of range expansion better than more cool-adapted historical occupants (Day et al., 2018). Adaptive evolution and plasticity may enable high population growth rates in newly-colonized areas (Angert et al., 2020; Usui et al., 2023), but this possibility can only be directly tested with long term population trend data. While some species experienced range retractions, these may result from sampling variability or stochastic population fluctuations along the northern range edge.

Increasing frequency and severity of extreme weather limited species’ geographical range responses (Table 2). This trend was independent of functional traits that are mechanistically linked to species’ climate change responses, such as dispersal ability or habitat preference. Extreme temperatures can reduce population sizes, leading to local extinctions (Román-Palacios and Wiens, 2020), and reducing the likelihood of range expansions (Mair et al., 2014). In odonates, experimental evidence has demonstrated that larval mortality rises with short-term extreme weather (McCauley et al., 2015). Individuals that shift phenologies earlier in the season to avoid climate extremes could still be exposed to harmful conditions (Iler et al., 2021); for example, odonate populations that respond to unusually warm spring temperatures may experience high mortality if temperatures return to seasonal conditions. Species that experience extreme conditions may then be unable to successfully shift in time, reducing population sizes, and reducing the likelihood of range shifts.

In contrast to previous work demonstrating that range and phenology shifts are at least partially determined by species traits (i.e. Sunday et al., 2015; Zografou et al., 2021), no functional trait, or combination of traits, explained these shifts in North American and European Odonata. While we could not capture all functional traits in this analysis, our results are consistent with other work that identifies climate velocity and sensitivity as the best predictors of range shifts and thermal preferences tracking in marine systems (Pinsky et al., 2013; Schuetz et al., 2019). Species’ tolerances to increasingly variable temperatures also helps to predict extinction risk during climate change (Kerr, 2020; Rocha-Ortega et al., 2020). The extent to which species’ traits actually determine rates of range and phenological shifts, rather than occasionally correlated with them, is worth considering further, but functional traits do not systematically drive patterns in these shifts among Odonates in North America and Europe.

The geographic positions of species’ ranges determine the local pressures and environmental factors to which they are exposed (MacLean and Beissinger, 2017; Pacifici et al., 2020), potentially masking or confounding the effects of traits that evolved under conditions determined by range geography (Schuetz et al., 2019). This process could cause trait-related trends to differ across levels of biological organization (Srivastava et al., 2021), from local populations (where traits might be critical) to biogeographical extents (where traits might be unrelated to range or phenological shifts; Grewe et al., 2013; Gutiérrez and Wilson, 2021; Sunday et al., 2015; Zografou et al., 2021).

Given that species’ functional traits did not predict temporal or geographic responses, it is unsurprising that species’ responses were also independent of phylogenetic history (Franke et al., 2022). The phylogenetic approach did not improve model predictions in any model that we tested, and there was no phylogenetic signal in either response according to Pagel’s lambda and Blomberg’s K (Table 2). These results are consistent with previous work that found no phylogenetic trend in local odonate population extinctions (Suhonen et al., 2022). There may be strong variation in thermal niches among closely related species: species that are geographically isolated adapt to different local climates, while species that co-occur may experience divergent selection within their climate tolerances (Schuetz et al., 2019).

It remains unclear if range and phenology shifts relate to trends in abundance, but our results suggest that there may be ‘winners’ and ‘losers’ under climate change (Figure 4). Climate ‘winners’, species that are shifting in space and time, may require more limited conservation intervention. Species expanding their ranges could be better supported if habitat area and connectivity are conserved, facilitating climate-driven range shifts (Littlefield et al., 2019). Species only shifting their phenologies may require further study, as phenology shifts may have positive or negative impacts on abundance (Iler et al., 2021). Climate ‘losers’, species that are failing to shift in both space and time, may require more direct conservation intervention, such as managed relocation (Richardson et al., 2009). Species that did not shift their ranges northwards or advance their phenology included *Coenagrion mercuriale*, a European species that is listed as near threatened by the IUCN Red List (IUCN, 2021), and is projected to lose 68% of its range by 2035 (Jaeschke et al., 2013). This group also includes *Coenagrion resolutum,* a common North American damselfly (Swaegers et al., 2014), for which we could not find evidence of decline. This may be due in part to the greater area of intact habitat available in North American compared to Europe, enabling *C. resolutum* to maintain larger populations that are less vulnerable to stochastic climate events. Still, this and other species failing to shift in range or phenology should be assessed for population health, as this species could be carrying an unobserved extinction debt. Our analysis of phenology and range shifts should be repeated in other taxa, as it may offer a method of identifying conservation actions among species groups.

**Figure 4:**
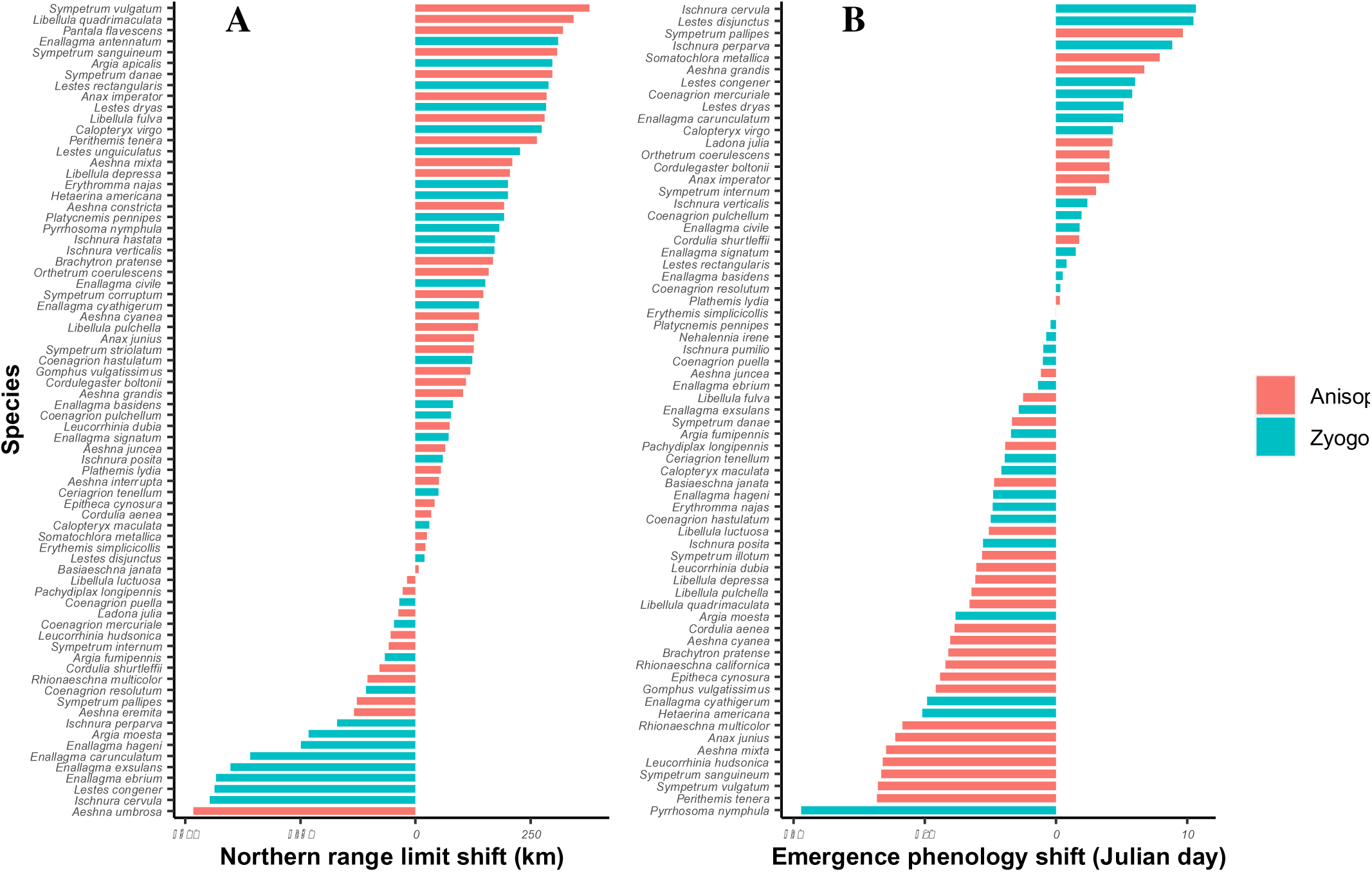
Distribution of Northern range limit shifts (Panel A, kilometers) and emergence phenology shift (Panel B, Julian day) of 76 European and North American odonate species between a recent time period (2008 - 2018) and a historical time period (1980 - 2002). Anisoptera (dragonflies) are shown in pink, Zygoptera (damselflies) are shown in blue.

Understanding how range and phenology shifts vary across species, and what drives this variation, is increasingly urgent as climate change alters local and regional environmental conditions. Here, we showed that odonate species exhibit convergent responses of range and phenology shifts across continents. While species with southern distributions were more likely to shift their ranges, increasing temperature variation limited geographical range responses among species in both Europe and North America. Climate change is associated with increasing variability as well as shifting mean conditions, contributing to species decline and even local extinction risks (Duffy et al., 2022). In this study, where species are found (i.e. their range geographies) determines whether they are exposed and respond to such negative pressures. Simultaneous consideration of shifts in range and phenology is a powerful and necessary approach to test aspects of species’ vulnerabilities to rapid global changes. By considering both the seasonal and range dynamics of species, emergent and convergent climate change responses across continents become clear for this well-studied group of predatory insects.

## Supplementary Information

### Odonate data

The full raw dataset includes ∼2 million Odonata occurrence records. We first removed inadequate records from our primary dataset. Inadequate records were identified as those with incomplete information for species identification, year, or locality, inaccurate georeferenced points, and duplicate records. We retained ∼1,100,000 unique species-location-date observations. There are 452,929 records in the first time period (1980 – 2002) and 641,590 observations in the second time period (2008 – 2018).

### Phenology and range position estimates

We have not tried to interpret noise in phenology shift estimates, which could occur due to sampling issues, or to another biologically meaningful eco-evolutionary response. Our methods do not enable tests of those ideas, however. For example, *Aeshna umbrosa* has a surprisingly high phenology shift (>25 days) but a very strong range retraction (<-300 km). For this species, we retained 8 quadrats to calculate mean phenology shift estimate. It is possible that a section of the range was lost in which emergence was especially early, compared to other regions, due to local adaptations. This pattern may affect results where species appear to shift emergence dates earlier/later, but phenology shifts are affected by parts of the range that remain occupied.

We put in place criteria to make sure to include species with range shifts likely to result from climate change effects, rather than from sampling issues or to other anthropogenic pressures such as land use change, or sudden land use intensification. *Nehalennia irene* was thus removed, as its range positions were highly unusual, and likely due to other factors than global change (>800 km). *Libellula quadrimaculata* was removed due to sampling intensity discrepancies between time periods, having over 10,000 extra points in the second period compared to the first.

Temporal and spatial bias are likely to be present in opportunistic data, but they are less likely to impact long term factors of species’ distributions if including data from as many sources as possible, and that span across large geographic and temporal scopes (Pyke and Ehrlich, 2010; Zattara and Aizen, 2021). Here, we put in place several criteria in careful interpretation of results, as described in the Methods section of the main text. Further, among preliminary examinations of the data, we test for the effect of sampling onto our phenology and range shift estimates. We found no effect of sampling on range of phenology responses that we calculated here. We tested a linear mixed model with species as a random effect that assesses whether sampling intensity change per quadrat (where sampling intensity corresponds to sampling per species/quadrat) predicts species’ phenology change per quadrat. The p-value of the predictor variable, sampling intensity change, was not significant. Secondly, we built a Generalized Linear Model to assess whether greater sampling overall (measured as the number of quadrats in which each species is sampled) predicts species’ range shifts that we computed. The p-value of the predictor variable, sampling change per species, was not significant. If spatial bias was present here, we may expect the number of quadrats in which species are found to increase the position of species’ northern range boundaries.

### Model information and statements

We report the full model statements below as executed with the *MCMCglmm* R package, including the priors, and the model of trait evolution to account for phylogeny. Prior structure followed standard practice according to continuous or binary response variable. The family “gaussian” or “threshold” was selected depending on the continuous or binary nature of the data, respectively. The number of iterations, burnin, and thinning parameters were tested and confirmed with a visual assessment of model convergence using the trace and density plots. It is common practice for the burnin parameter to correspond to 10% of the number of iterations.

### Model 1

Aphylo ← vcv(phylogeny, model = “Brownian”, corr = T)

Ainv ← inverseA(phylogeny, nodes = “TIPS”, scale = F)$Ainv

prior ← list(R=list(V=1,nu=0.002))

MCMCglmm(Phenological shifts ∼ Range shifts + Continent, ginverse = list(species = Ainv), family = “gaussian”, prior = prior, DIC = T, nitt = 150,000, thin = 100, burnin = 15,000, data)

### Model 2

Aphylo ← vcv(phylogeny, model = “Brownian”, corr = T)

Ainv ← inverseA(phylogeny, nodes = “TIPS”, scale = F)$Ainv

prior ← list(R=list(V=1,nu=0.002, fix = 1))

MCMCglmm(Range shifts ∼ Breeding habitat type + Distribution region + Flight duration + Egg laying habitat + Body size + Temperature variability, ginverse = list(species = Ainv), family = “threshold”, prior = prior, DIC = T, nitt = 150,000, thin = 100, burnin = 15,000, data)

### Model 3

Aphylo ← vcv(phylogeny, model = “Brownian”, corr = T)

Ainv ← inverseA(phylogeny, nodes = “TIPS”, scale = F)$Ainv

prior ← list(R=list(V=1,nu=0.002))

MCMCglmm(Phenology shift ∼ Breeding habitat type + Distribution region + Flight duration + Egg laying habitat + Body size + Temperature variability, ginverse = list(species = Ainv), family = “gaussian”, prior = prior, DIC = T, nitt = 150,000, thin = 100, burnin = 15,000, data)

**Table S1:** Samples of 76 North American and European odonate species from between 1980 and 2018 followed our criteria for quality observation records for inclusion in our analysis of geographical shifts. Species Northern Range Limits (NRL) are shown in this table, as well as range limit shifts. All range limit values are shown in kilometers from the equ10ator. We used the 10 most northern points of sampling in each time period to identify species’ NRL, as detailed in the Methods section of the main text.

| Species | Continent | NRL<br>(1980 -<br>2002) | NRL<br>(2008 -<br>2018) | NRL<br>shift |
| --- | --- | --- | --- | --- |
| *Aeshna constricta* | North-America | 5500.81 | 5692.6 | 191.79 |
| *Aeshna cyanea* | Europe | 6880.39 | 7018.26 | 137.87 |
| *Aeshna eremita* | North-America | 7527.81 | 7394.66 | -133.16 |
| *Aeshna grandis* | Europe | 7504.84 | 7607.68 | 102.83 |
| *Aeshna interrupta* | North-America | 7235.65 | 7286.83 | 51.18 |
| *Aeshna juncea* | Europe | 7747.56 | 7812.16 | 64.6 |
| *Aeshna mixta* | Europe | 6566.18 | 6776.3 | 210.12 |
| *Aeshna umbrosa* | North-America | 6691.19 | 6209.63 | -481.55 |
| *Anax imperator* | Europe | 6263.92 | 6548.57 | 284.65 |
| *Anax junius* | North-America | 5539.31 | 5666.11 | 126.8 |
| *Argia apicalis* | North-America | 4922.17 | 5219.38 | 297.22 |
| *Argia fumipennis* | North-America | 5235.2 | 5168.12 | -67.08 |
| *Argia moesta* | North-America | 5533.74 | 5301.98 | -231.77 |
| *Basiaeschna janata* | North-America | 5649.43 | 5655.77 | 6.34 |
| *Brachytron pratense* | Europe | 6737.42 | 6905.46 | 168.04 |
| *Calopteryx maculata* | North-America | 5329.56 | 5359.13 | 29.57 |
| *Calopteryx virgo* | Europe | 7275.9 | 7549.35 | 273.45 |
| *Ceriagrion tenellum* | Europe | 5900.55 | 5950.63 | 50.08 |
| *Coenagrion hastulatum* | Europe | 7528.94 | 7651.51 | 122.57 |
| *Coenagrion mercuriale* | Europe | 5913.57 | 5867.06 | -46.52 |
| *Coenagrion puella* | Europe | 6904.7 | 6869.58 | -35.12 |
| *Coenagrion pulchellum* | Europe | 7076.45 | 7152.81 | 76.36 |
| *Coenagrion resolutum* | North-America | 7516.44 | 7408.7 | -107.73 |
| *Cordulegaster boltonii* | Europe | 7237.43 | 7346.93 | 109.49 |
| *Cordulia aenea* | Europe | 7400.12 | 7434.02 | 33.9 |
| *Cordulia shurtleffii* | North-America | 7499.82 | 7421.98 | -77.84 |
| *Enallagma antennatum* | North-America | 5037.73 | 5347.01 | 309.28 |
| *Enallagma basidens* | North-America | 4768.61 | 4850.05 | 81.44 |
| *Enallagma carunculatum* | North-America | 5950.86 | 5592.47 | -358.38 |
| *Enallagma civile* | North-America | 5322.59 | 5473.78 | 151.19 |
| *Enallagma cyathigerum* | Europe | 7594.97 | 7733.06 | 138.09 |
| *Enallagma ebrium* | North-America | 6341.31 | 5907.69 | -433.62 |
| *Enallagma exsulans* | North-America | 5655.94 | 5253.94 | -402 |
| *Enallagma hageni* | North-America | 6113.97 | 5865.52 | -248.45 |
| *Enallagma signatum* | North-America | 5208.92 | 5281.09 | 72.17 |
| *Epitheca cynosura* | North-America | 5541.19 | 5582.56 | 41.37 |
| *Erythemis simplicicollis* | North-America | 5271 | 5292.65 | 21.65 |
| *Erythromma najas* | Europe | 7171.87 | 7372.13 | 200.25 |
| *Gomphus vulgatissimus* | Europe | 6890.76 | 7010.12 | 119.37 |
| *Hetaerina americana* | North-America | 5044.41 | 5244.49 | 200.08 |
| *Ischnura cervula* | North-America | 5973.57 | 5526.83 | -446.74 |
| *Ischnura hastata* | North-America | 4725.07 | 4897.71 | 172.65 |
| *Ischnura perparva* | North-America | 5608.92 | 5438.53 | -170.4 |
| *Ischnura posita* | North-America | 5084.08 | 5143.79 | 59.71 |
| *Ischnura pumilio* | Europe | 6213.77 | 6769.62 | 555.85 |
| *Ischnura verticalis* | North-America | 5579.68 | 5750.9 | 171.22 |
| *Ladona julia* | North-America | 5908.68 | 5871.84 | -36.83 |
| *Lestes congener* | North-America | 6240.48 | 5803.75 | -436.74 |
| *Lestes disjunctus* | North-America | 7294.75 | 7314.26 | 19.51 |
| *Lestes dryas* | Europe | 6824.89 | 7108.33 | 283.44 |
| *Lestes rectangularis* | North-America | 5263.85 | 5552.64 | 288.79 |
| *Lestes unguiculatus* | North-America | 5803.91 | 6030.62 | 226.71 |
| *Leucorrhinia dubia* | Europe | 7675.72 | 7749.01 | 73.29 |
| *Leucorrhinia hudsonica* | North-America | 7500.27 | 7446.85 | -53.42 |
| *Libellula depressa* | Europe | 6810.85 | 7015.37 | 204.52 |
| *Libellula fulva* | Europe | 6602.18 | 6882.5 | 280.32 |
| *Libellula luctuosa* | North-America | 5337.86 | 5320.1 | -17.76 |
| *Libellula pulchella* | North-America | 5556.31 | 5692.14 | 135.83 |
| *Orthetrum coerulescens* | Europe | 6744.55 | 6903.01 | 158.46 |
| *Pachydiplax longipennis* | North-America | 5495.02 | 5467.11 | -27.92 |
| *Pantala flavescens* | North-America | 5190.25 | 5510.15 | 319.9 |
| *Perithemis tenera* | North-America | 4929.38 | 5192.47 | 263.09 |
| *Plathemis lydia* | North-America | 5539.41 | 5594.8 | 55.38 |
| *Platycnemis pennipes* | Europe | 6934.22 | 7125.72 | 191.5 |
| *Pyrrosoma nymphula* | Europe | 7131.07 | 7313.37 | 182.3 |
| *Rhionaeschna californica* | North-America | 5641.53 | 5458.21 | -183.32 |
| *Rhionaeschna multicolor* | North-America | 5581 | 5477.34 | -103.66 |
| *Somatochlora metallica* | Europe | 7718.67 | 7743.21 | 24.54 |
| *Somatochlora*<br>*semicircularis* | North-America | 6533.02 | 5994.13 | -538.89 |
| *Sympetrum corruptum* | North-America | 5585.68 | 5733.2 | 147.52 |
| *Sympetrum danae* | Europe | 7268.54 | 7565.29 | 296.75 |
| *Sympetrum internum* | North-America | 7179.45 | 7121.38 | -58.07 |
| *Sympetrum pallipes* | North-America | 5655.65 | 5527.84 | -127.8 |
| *Sympetrum sanguineum* | Europe | 6663.43 | 6970.56 | 307.13 |
| *Sympetrum striolatum* | Europe | 6972.22 | 7098.74 | 126.52 |
| *Sympetrum vulgatum* | Europe | 6897.33 | 7274.78 | 377.45 |

**Table S2:** 66 species sampled across North America and Europe between 1980 and 2018 followed our criteria for quality observation records for inclusion in our analysis of emergence phenology shifts. Mean phenological shifts (PS) is measured in the number of Julian days comparing both time periods, as estimated using the Weibull distribution (See Methods). We also report the number of 200 X 200 quadrats used to calculate phenology estimates per species.

| Species | Number of quadrats | Mean PS |
| --- | --- | --- |
| *Aeshna cyanea* | 37 | -8.05 |
| *Aeshna grandis* | 43 | 6.73 |
| *Aeshna juncea* | 34 | -1.11 |
| *Aeshna mixta* | 22 | -12.95 |
| *Aeshna umbrosa* | 8 | 27.42 |
| *Anax imperator* | 27 | 4.05 |
| *Anax junius* | 19 | -12.21 |
| *Argia fumipennis* | 8 | -3.39 |
| *Argia moesta* | 11 | -7.62 |
| *Basiaeschna janata* | 2 | -4.69 |
| *Brachytron pratense* | 24 | -8.18 |
| *Calopteryx maculata* | 15 | -4.14 |
| *Calopteryx virgo* | 41 | 4.35 |
| *Ceriagrion tenellum* | 14 | -3.88 |
| *Coenagrion hastulatum* | 20 | -4.97 |
| *Coenagrion mercuriale* | 11 | 5.80 |
| *Coenagrion puella* | 33 | -1.00 |
| *Coenagrion pulchellum* | 29 | 1.94 |
| *Coenagrion resolutum* | 2 | 0.33 |
| *Cordulegaster boltonii* | 34 | 4.08 |
| *Cordulia aenea* | 31 | -7.72 |
| *Cordulia shurtleffii* | 4 | 1.78 |
| *Enallagma basidens* | 2 | 0.52 |
| *Enallagma carunculatum* | 8 | 5.10 |
| *Enallagma civile* | 11 | 1.82 |
| *Enallagma cyathigerum* | 46 | -9.81 |
| *Enallagma ebrium* | 7 | -1.34 |
| *Enallagma exsulans* | 8 | -2.83 |
| *Enallagma hageni* | 5 | -4.79 |
| *Enallagma signatum* | 6 | 1.51 |
| *Epitheca cynosura* | 4 | -8.82 |
| *Erythemis simplicicollis* | 15 | 0.01 |
| *Erythromma najas* | 35 | -4.82 |
| *Gomphus vulgatissimus* | 12 | -9.14 |
| *Hetaerina americana* | 6 | -10.15 |
| *Ischnura cervula* | 3 | 10.65 |
| *Ischnura perparva* | 2 | 8.85 |
| *Ischnura posita* | 12 | -5.55 |
| *Ischnura pumilio* | 14 | -0.97 |
| *Ischnura verticalis* | 16 | 2.39 |
| *Ladona julia* | 9 | 4.29 |
| *Lestes congener* | 9 | 6.03 |
| *Lestes disjunctus* | 4 | 10.48 |
| *Lestes dryas* | 6 | 5.13 |
| *Lestes rectangularis* | 12 | 0.81 |
| *Leucorrhinia dubia* | 18 | -6.05 |
| *Leucorrhinia hudsonica* | 4 | -13.17 |
| *Libellula depressa* | 30 | -6.13 |
| *Libellula fulva* | 16 | -2.50 |
| *Libellula luctuosa* | 15 | -5.11 |
| *Libellula pulchella* | 14 | -6.43 |
| *Orthetrum coerulescens* | 22 | 4.11 |
| *Pachydiplax longipennis* | 13 | -3.84 |
| *Perithemis tenera* | 5 | -13.60 |
| *Plathemis lydia* | 14 | 0.30 |
| *Platycnemis pennipes* | 30 | -0.40 |
| *Pyrrhosoma nymphula* | 39 | -19.40 |
| *Rhionaeschna californica* | 2 | -8.41 |
| *Rhionaeschna multicolor* | 4 | -11.68 |
| *Somatochlora metallica* | 31 | 7.91 |
| *Sympetrum danae* | 32 | -3.34 |
| *Sympetrum internum* | 5 | 3.05 |
| *Sympetrum pallipes* | 3 | 9.67 |
| *Sympetrum sanguineum* | 28 | -13.30 |
| *Sympetrum vulgatum* | 16 | -13.56 |

**Table S3:** Ecological and geographical traits of 76 North American and European odonate species used in this work. Field guides (Cannings, 2002; Jones et al., 2008; Paulson, 2012) and existing trait databases (Powney et al., 2014; Waller et al., 2019) were used to build this dataset. Habitat type represents species’ breeding habitat, and can have a value of lentic, lotic, or both types. Distribution shows the general geographic position of each species’ range, which can be widespread (W), southern (S), northern (N), southern and widespread (SW), or northern and widespread (NW). Oviposition type corresponds to egg laying inside plants (endophytic) as opposed to directly in water or on plants (exophytic). Body size is measured as body length in mm. In the case that body length was given as a maximum and minimum value, we used the

| Species | Habitat | Distribution | Flight | Oviposition | Body size |
| --- | --- | --- | --- | --- | --- |
| *Aeshna constricta* | Both | W | 2.5 | Endophytic | 69 |
| *Aeshna cyanea* | Lentic | S | 3 | Endophytic | 71.5 |
| *Aeshna eremita* | Both | N-W | 3 | Endophytic | 72.5 |
| *Aeshna grandis* | Both | N | 4 | Endophytic | 73.5 |
| *Aeshna interrupta* | Both | W | 3 | Endophytic | 66.5 |
| *Aeshna juncea* | Lentic | N | 4.5 | Endophytic | 75.5 |
| *Aeshna mixta* | Both | S | 2.5 | Endophytic | 60 |
| *Aeshna umbrosa* | Both | W | 3 | Endophytic | 68.5 |
| *Anax imperator* | Lentic | S | 2.5 | Endophytic | 76.5 |
| *Anax junius* | Both | S-W | 7 | Endophytic | 74 |
| *Argia apicalis* | Both | S-W | 4 | Endophytic | 36.5 |
| *Argia fumipennis* | Both | S-W | 3.5 | Endophytic | 31.5 |
| *Argia moesta* | Lotic | S-W | 3 | Endophytic | 39.5 |
| *Basiaeschna janata* | Both | W | 1 | Endophytic | 59 |
| *Brachytron pratense* | Lentic | S | 2 | Endophytic | 58.5 |
| *Calopteryx maculata* | Lotic | S-W | 3 | Endophytic | 48 |
| *Calopteryx virgo* | Lotic | W | 3.5 | Endophytic | 47 |
| *Ceriagrion tenellum* | Both | S | 3 | Endophytic | 30 |
| *Coenagrion hastulatum* | Lentic | N | 2.5 | Endophytic | 32 |
| *Coenagrion mercuriale* | Lotic | S | 2.5 | Endophytic | 29.5 |
| *Coenagrion puella* | Lentic | S | 3.5 | Endophytic | 34 |
| *Coenagrion pulchellum* | Both | S | 2 | Endophytic | 36 |
| *Coenagrion resolutum* | Lentic | N-W | 2.5 | Endophytic | 30 |
| *Cordulegaster boltonii* | Lotic | W | 3.5 | Endophytic | 77 |
| *Cordulia aenea* | Lentic | S | 2 | Exophytic | 51 |
| *Cordulia shurtleffii* | Both | N-W | 2 | Exophytic | 46 |
| *Enallagma antennatum* | Both | S-W | 2 | Endophytic | 30 |
| *Enallagma basidens* | Both | S-W | 4.5 | Endophytic | 24.5 |
| *Enallagma carunculatum* | Both | W | 3.5 | Endophytic | 31.5 |
| *Enallagma civile* | Both | S-W | 3.5 | Endophytic | 33.5 |
| *Enallagma cyathigerum* | Both | W | 3.5 | Endophytic | 32 |
| *Enallagma ebrium* | Both | N-W | 2.5 | Endophytic | 31 |
| *Enallagma exsulans* | Both | S-W | 3.5 | Endophytic | 34 |
| *Enallagma hageni* | Both | N-W | 2.5 | Endophytic | 30 |
| *Enallagma signatum* | Both | S-W | 2.5 | Endophytic | 32.5 |
| *Epitheca cynosura* | Both | S-W | 2.5 | Endophytic | 40.5 |
| *Erythemis simplicicollis* | Both | S-W | 2.5 | Endophytic | 41 |
| *Erythromma najas* | Both | S | 3 | Endophytic | 33 |
| *Gomphus vulgatissimus* | Lotic | S | 2 | Exophytic | 47.5 |
| *Hetaerina americana* | Lotic | S-W | 5 | Endophytic | 42 |
| *Ischnura cervula* | Both | W | 6 | Endophytic | 27.5 |
| *Ischnura hastata* | Lentic | S-W | 3.5 | Endophytic | 24 |
| *Ischnura perparva* | Both | S-W | 5 | Endophytic | 26.5 |
| *Ischnura posita* | Both | S-W | 3.5 | Endophytic | 25 |
| *Ischnura pumilio* | Lentic | S | 3 | Endophytic | 29 |
| *Ischnura verticalis* | Both | W | 3.5 | Endophytic | 26.5 |
| *Ladona julia* | Lentic | N-W | 2 | Endophytic | 41.5 |
| *Lestes congener* | Lentic | W | 2.5 | Endophytic | 36.5 |
| *Lestes disjunctus* | Both | W | 2.5 | Endophytic | 37.5 |
| *Lestes dryas* | Lentic | S | 2 | Endophytic | 37.5 |
| *Lestes rectangularis* | Both | S-W | 2.5 | Endophytic | 45 |
| *Lestes unguiculatus* | Both | W | 3 | Endophytic | 37.5 |
| *Leucorrhinia dubia* | Lentic | N | 2.5 | Exophytic | 33.5 |
| *Leucorrhinia hudsonica* | Lentic | N-W | 2 | Endophytic | 29.5 |
| *Libellula depressa* | Lentic | S | 2.5 | Exophytic | 43.5 |
| *Libellula fulva* | Lentic | S | 2 | Exophytic | 43.5 |
| *Libellula luctuosa* | Lentic | S-W | 2.5 | Exophytic | 46 |
| *Libellula pulchella* | Both | W | 2.5 | Endophytic | 54.5 |
| *Orthetrum coerulescens* | Lotic | S | 3 | Exophytic | 40.5 |
| *Pachydiplax longipennis* | Both | S-W | 3 | Endophytic | 35.5 |
| *Pantala flavescens* | Both | S | 3 | Exophytic | 48.5 |
| *Perithemis tenera* | Both | S-W | 4.5 | Exophytic | 22.5 |
| *Plathemis lydia* | Both | S-W | 3 | Exophytic | 45 |
| *Platycnemis pennipes* | Lotic | S | 2.5 | Endophytic | 36 |
| *Pyrrhosoma nymphula* | Lentic | W | 1.5 | Endophytic | 34.5 |
| *Rhionaeschna californica* | Lentic | S-W | 4 | Endophytic | 60.5 |
| *Rhionaeschna multicolor* | Both | S-W | 2 | Endophytic | 67 |
| *Somatochlora metallica* | Both | W | 1.5 | Exophytic | 53 |
| *Somatochlora*<br>*semicircularis* | Lentic | W | 2 | Exophytic | 49.5 |
| *Sympetrum corruptum* | Both | S-W | 5 | Exophytic | 40.5 |
| *Sympetrum danae* | Lentic | W | 2 | Exophytic | 31.5 |
| *Sympetrum internum* | Lentic | W | 4 | Exophytic | 33.5 |
| *Sympetrum pallipes* | Both | S-W | 2 | Endophytic | 36 |
| *Sympetrum sanguineum* | Lentic | S | 3.5 | Exophytic | 36.5 |
| *Sympetrum striolatum* | Both | W | 5 | Exophytic | 39.5 |
| *Sympetrum vulgatum* | Lentic | S | 2 | Exophytic | 37.5 |

**Table S4:** Credible intervals of all MCMCglmm models testing predictions regarding the range and phenology shifts across 66 odonate species in North America and Europe. These models are detailed in *Model information and statements* of the Supplementary Information.

|  | Lower credible interval | Upper credible interval |
| --- | --- | --- |
| **Phenological shifts ~ Range shifts** |  |  |
| (Intercept) | -0.26 | 0.49 |
| Range shifts | -0.71 | -0.16 |
| Continent | -0.75 | 0.30 |
| **Range shifts ~ Range geography + T ° variability** |  |  |
| (intercept) | -1.22 | -0.12 |
| Southern range | 0.36 | 1.56 |
| Widespread | -0.32 | 1.08 |
| T ° variability | -0.60 | -0.18 |

**Figure S1:**
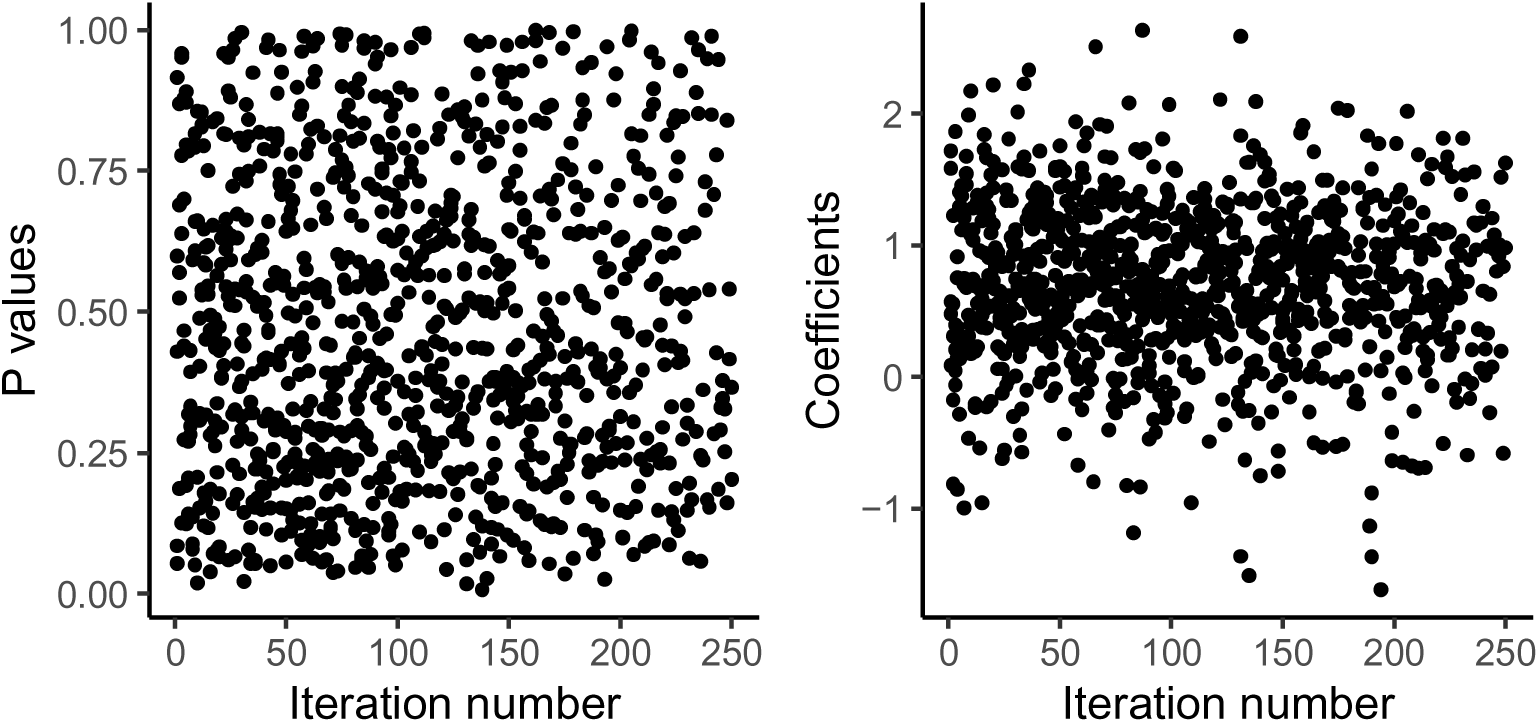
P-values and coefficients of 1000 GLM iterations testing whether range shifts as calculated from random datasets predict the range shifts measured in the study. Each point shows the results of a single GLM model, with measured range shifts as the dependant variable and randomized range shifts as the independent variable.

**Figure S2:**
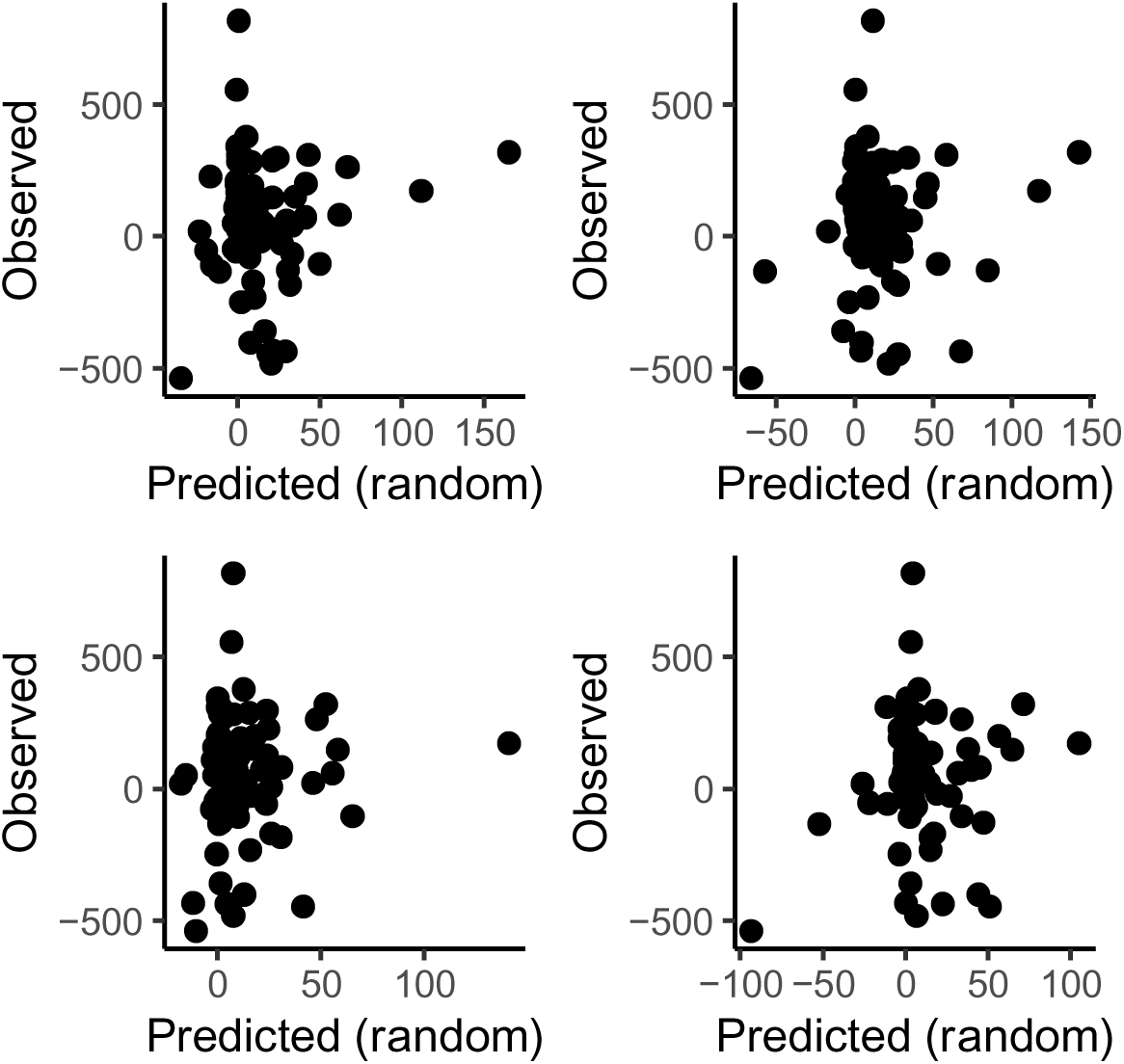
Observed range shifts in km from the equator, against randomized predicted values according to 4 random datasets. Points represent species and each pane contains a different set of random data in calculations of randomized range shifts. There is no consistent relationship among 1000 iterations.

**Figure S3:**
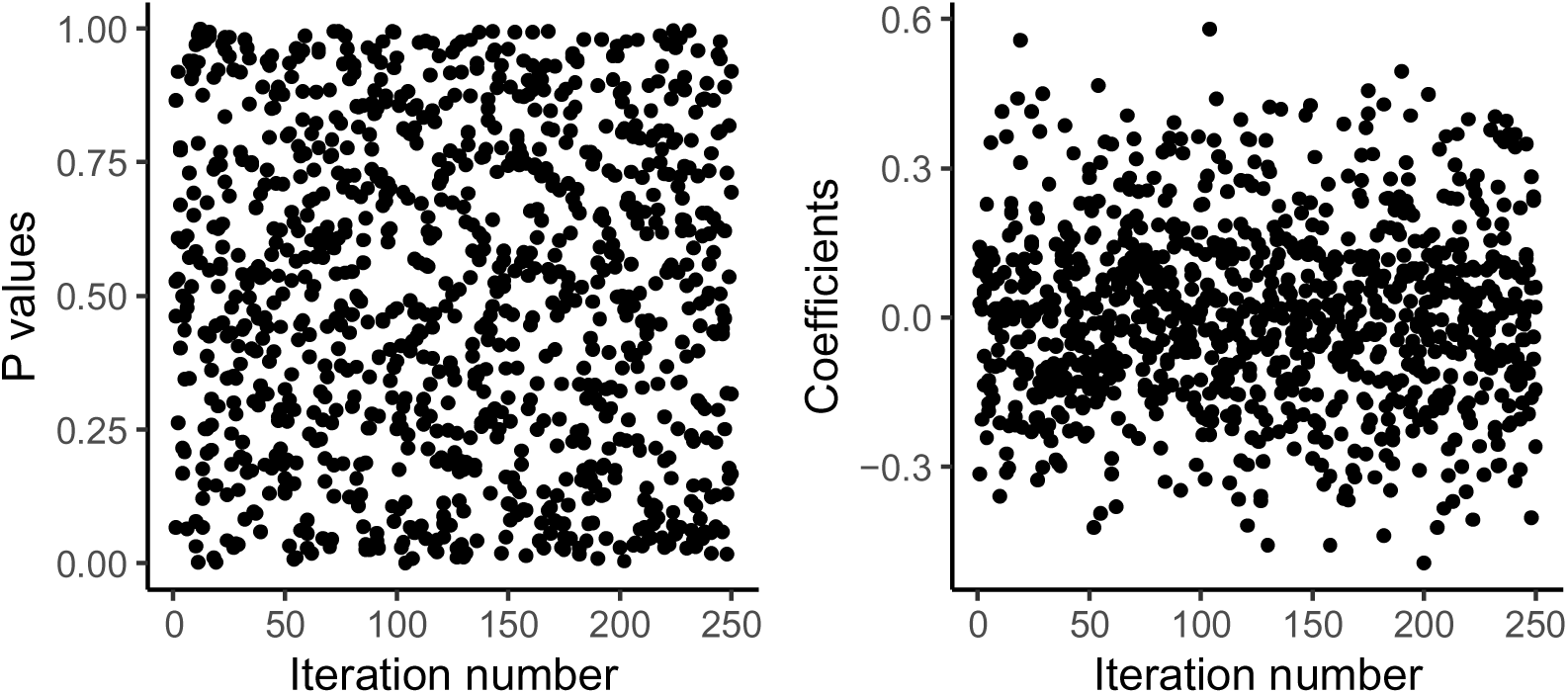
P-values and coefficients of 1000 GLM iterations testing whether phenology shifts as calculated from random datasets predict the phenology shifts measured in the study. Each point shows the results of a single GLM model, with measured phenology shifts as the dependant variable and randomized phenology shifts as the independent variable.

**Figure S4:**
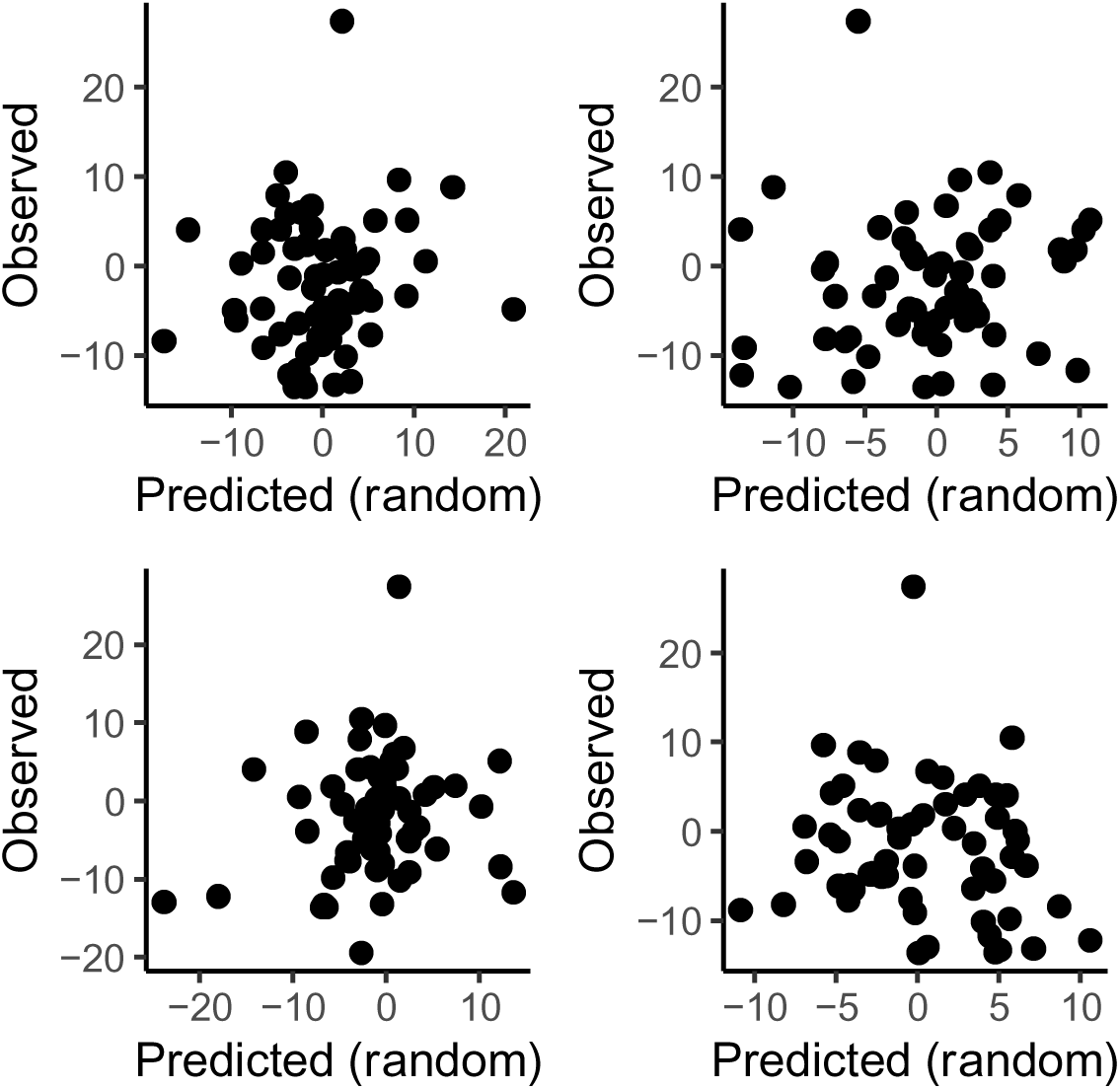
Observed phenology shifts in Julian day, against randomized predicted values according to 4 random datasets. Points represent species and each pane contains a different set of random data in calculations of randomized phenology shifts. There is no consistent relationship among 1000 iterations.

**Figure S7:**
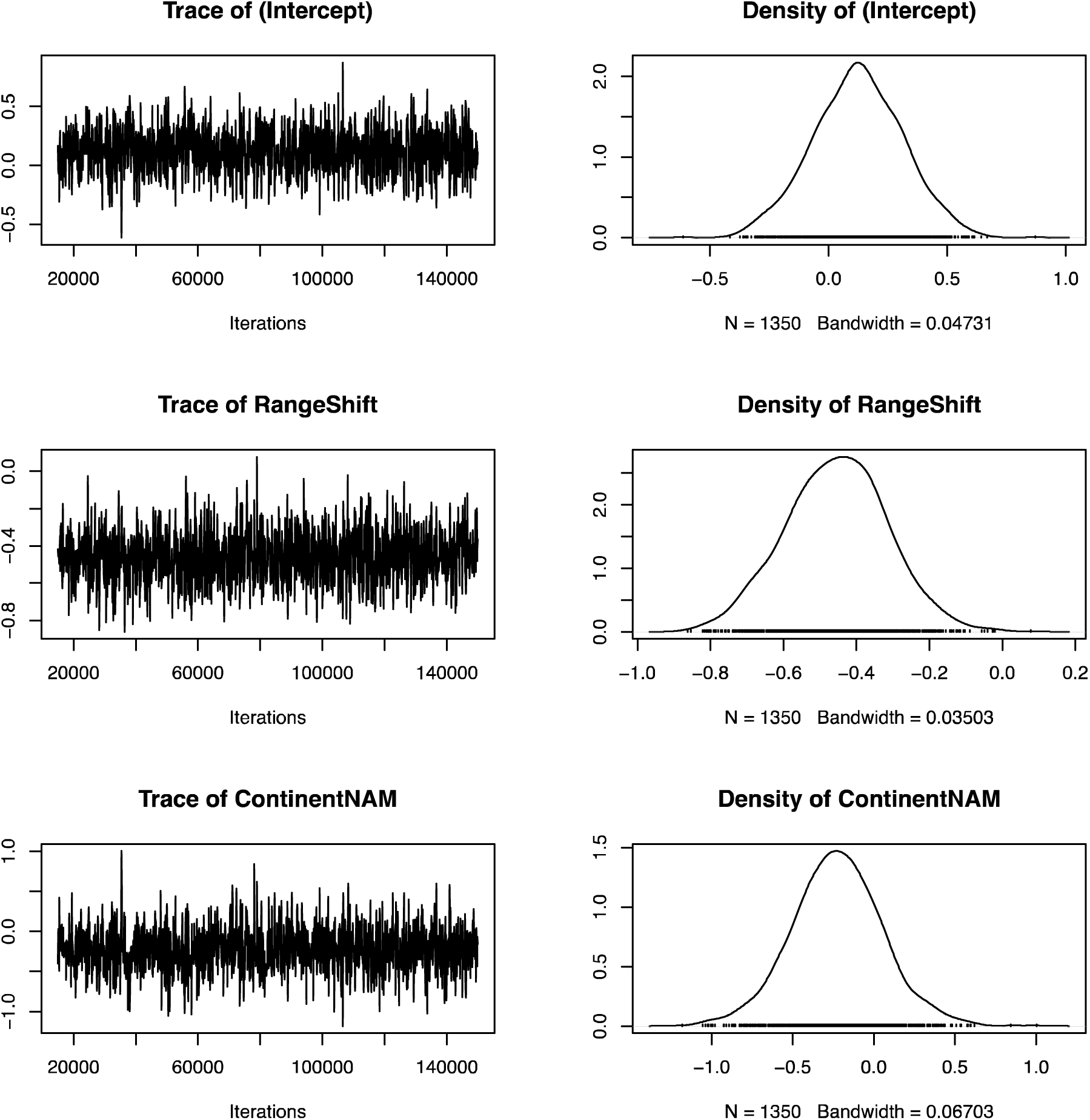
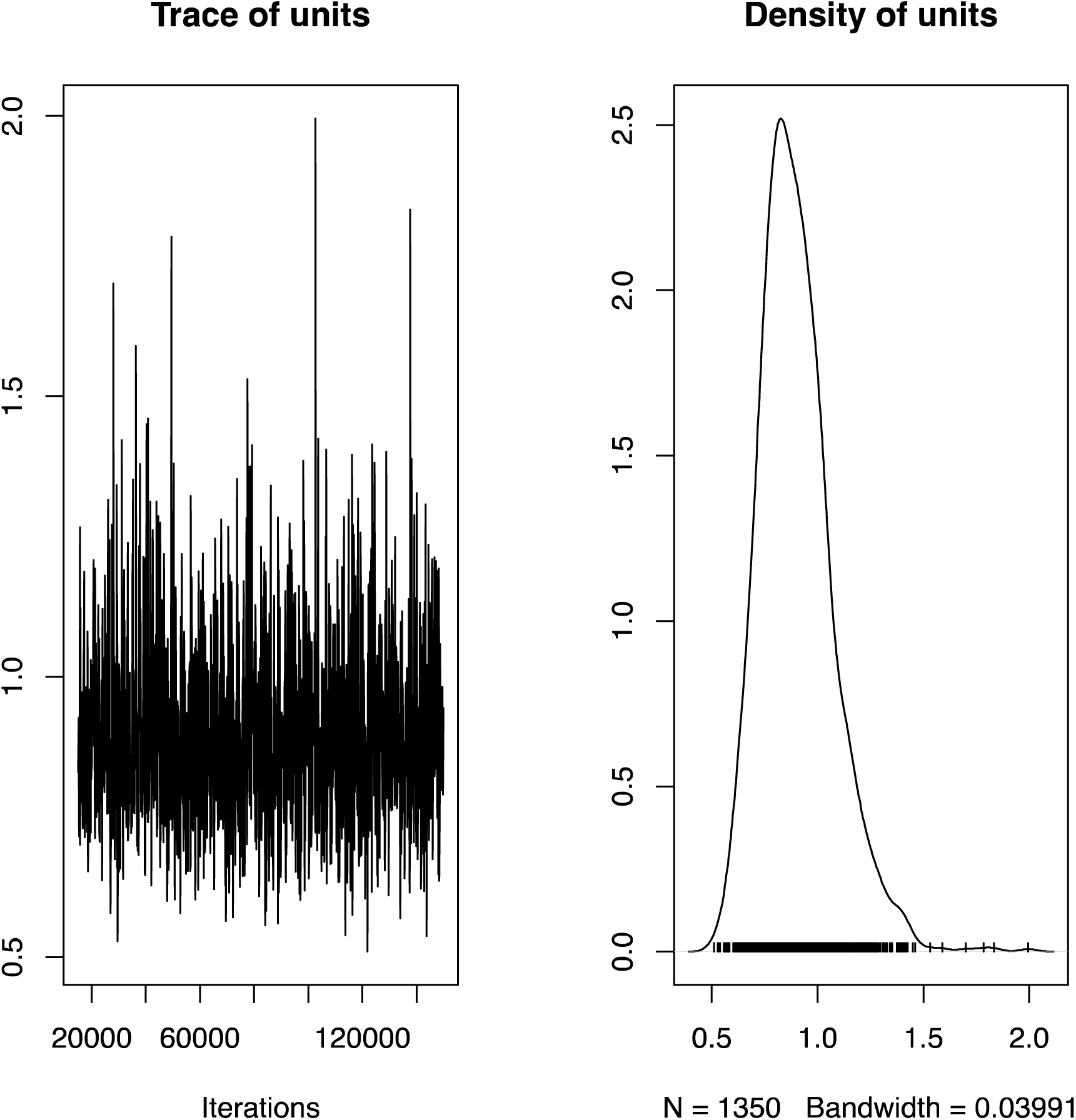
Panels A and B show the trace and density estimates of a phylogenetic mixed effects model exploring the relationship between range and phenology shifts in North American and European odonates (N = 66). 150,000 iterations were run to produce these results. These plots verify model convergence and absence of autocorrelation within the explanatory variables.

**Figure S8:**
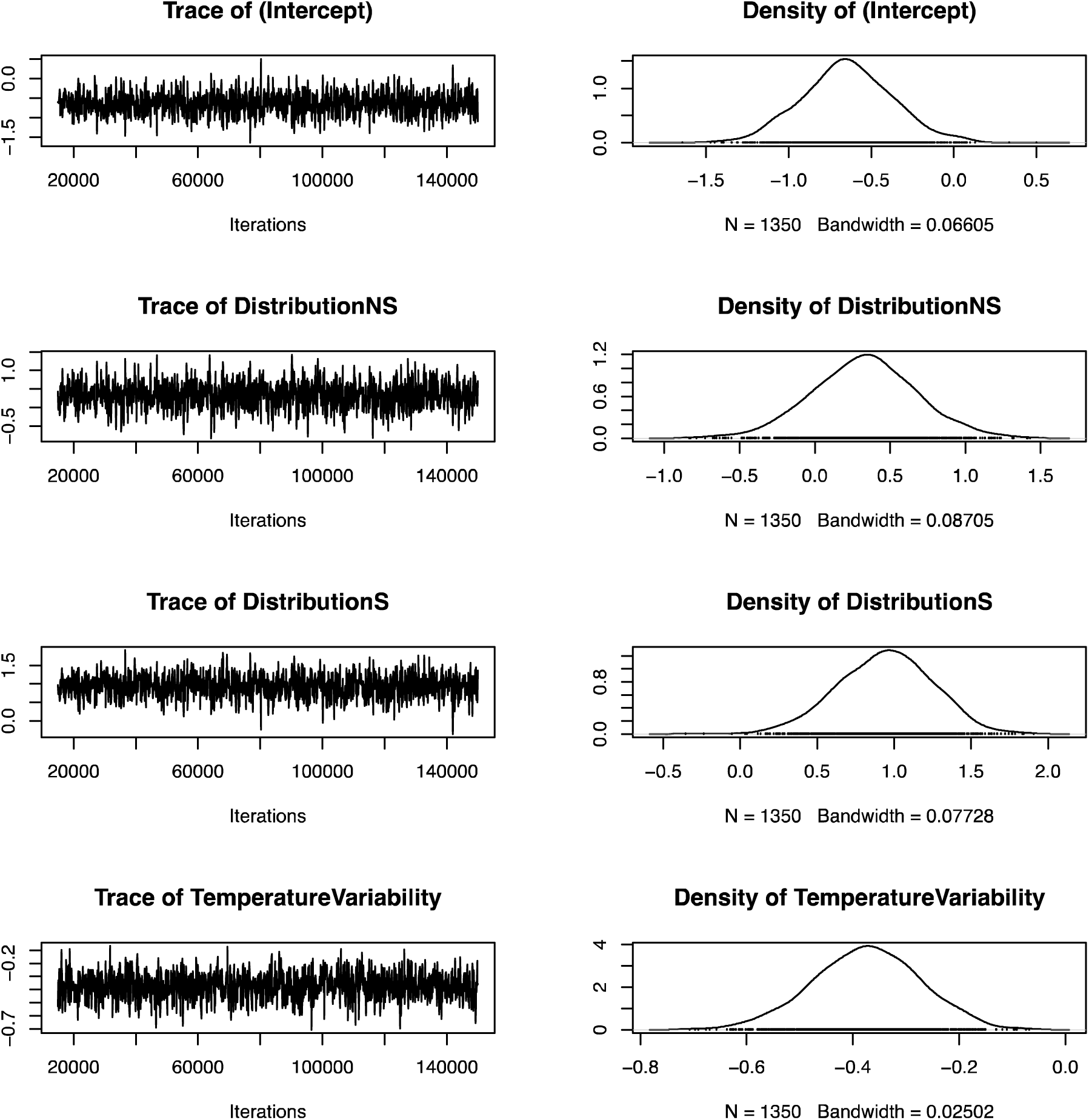
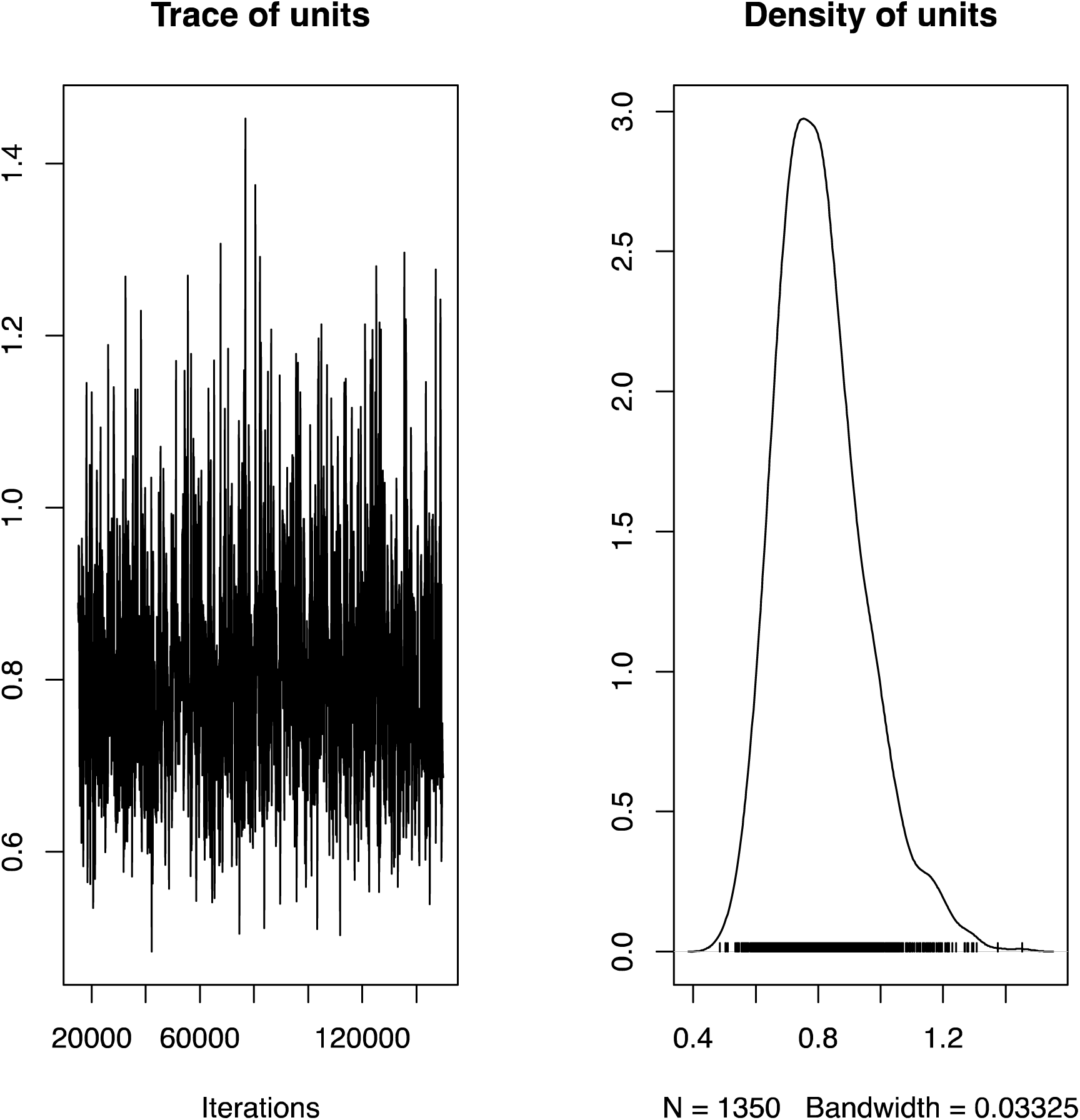
Panels A and B show the trace and density estimates a phylogenetic mixed effects model testing whether ecological traits, and geographic and climatic attributes predict range shifts in North American and European odonates (N = 76). 150,000 iterations were run to produce these results. Results of the best model, according to DIC, are shown here. These plots verify model convergence and absence of autocorrelation within the explanatory variables.

## References

Amano T, Freckleton RP, Queenborough SA, Doxford SW, Smithers RJ, Sparks TH, Sutherland WJ. 2014. Links between plant species’ spatial and temporal responses to a warming climate. Proceedings of the Royal Society B: Biological Sciences 281:20133017. doi:10.1098/rspb.2013.3017

Angert AL, Bontrager MG, Ågren J. 2020. What Do We Really Know About Adaptation at Range Edges? Annual Review of Ecology, Evolution, and Systematics 51:341–361. doi:10.1146/annurev-ecolsys-012120-091002

Angert AL, Crozier LG, Rissler LJ, Gilman SE, Tewksbury JJ, Chunco AJ. 2011. Do species’ traits predict recent shifts at expanding range edges? Ecology Letters 14:677–689. doi:10.1111/j.1461-0248.2011.01620.x

Ball-Damerow JE, M’Gonigle LK, Resh VH. 2014. Changes in occurrence, richness, and biological traits of dragonflies and damselflies (Odonata) in California and Nevada over the past century. Biodivers Conserv 23:2107–2126. doi:10.1007/s10531-014-0707-5

Ball-Damerow JE, Oboyski PT, Resh VH. 2015. California dragonfly and damselfly (Odonata) database: temporal and spatial distribution of species records collected over the past century. Zookeys 67–89. doi:10.3897/zookeys.482.8453

Bartomeus I, Stavert JR, Ward D, Aguado O. 2018. Historical collections as a tool for assessing the global pollination crisis. Philosophical Transactions of the Royal Society B: Biological Sciences 374:20170389. doi:10.1098/rstb.2017.0389

Beissinger SR, Riddell EA. 2021. Why Are Species’ Traits Weak Predictors of Range Shifts? Annual Review of Ecology, Evolution, and Systematics 52:47–66. doi:10.1146/annurev-ecolsys-012021-092849

Bowler DE, Eichenberg D, Conze K-J, Suhling F, Baumann K, Benken T, Bönsel A, Bittner T, Drews A, Günther A, Isaac NJB, Petzold F, Seyring M, Spengler T, Trockur B, Willigalla C, Bruelheide H, Jansen F, Bonn A. 2021. Winners and losers over 35 years of dragonfly and damselfly distributional change in Germany. Diversity and Distributions 27:1353–1366. doi:10.1111/ddi.13274

Bried JT, Rocha-Ortega M. 2023. Using range size to augment regional priority listing of charismatic insects. Biological Conservation 283:110098. doi:10.1016/j.biocon.2023.110098

Brooks SJ, Self A, Toloni F, Sparks T. 2014. Natural history museum collections provide information on phenological change in British butterflies since the late-nineteenth century. Int J Biometeorol 58:1749–1758. doi:10.1007/s00484-013-0780-6

Buckley LB, Kingsolver JG. 2012. Functional and Phylogenetic Approaches to Forecasting Species’ Responses to Climate Change. Annual Review of Ecology, Evolution, and Systematics 43:205–226. doi:10.1146/annurev-ecolsys-110411-160516

Cannings RA. 2002. Introducing the Dragonflies of British Columbia and Yukon. Royal British Columbia Museum.

Cardillo M, Mace GM, Gittleman JL, Jones KE, Bielby J, Purvis A. 2008. The predictability of extinction: biological and external correlates of decline in mammals. Proceedings of the Royal Society B: Biological Sciences 275:1441–1448. doi:10.1098/rspb.2008.0179

Chen I-C, Hill JK, Ohlemüller R, Roy DB, Thomas CD. 2011. Rapid Range Shifts of Species Associated with High Levels of Climate Warming. Science 333:1024–1026.

Chichorro F, Juslén A, Cardoso P. 2019. A review of the relation between species traits and extinction risk. Biological Conservation 237:220–229. doi:10.1016/j.biocon.2019.07.001

Conrad KF, Willson KH, Harvey IF, Thomas CJ, Sherratt TN. 1999. Dispersal characteristics of seven odonate species in an agricultural landscape. Ecography 22:524–531. doi:10.1111/j.1600-0587.1999.tb01282.x

Cooper N, Bielby J, Thomas GH, Purvis A. 2008. Macroecology and extinction risk correlates of frogs. Global Ecology and Biogeography 17:211–221. doi:10.1111/j.1466-8238.2007.00355.x

Córdoba-Aguilar A, Beatty C, Bried JT. 2023. Dragonflies and Damselflies: Model Organisms for Ecological and Evolutionary Research. Oxford University Press.

Davis MB, Shaw RG, Etterson JR. 2005. Evolutionary responses to changing climate. Ecology 86:1704–1714. doi:10.1890/03-0788

Day PB, Stuart-Smith RD, Edgar GJ, Bates AE. 2018. Species’ thermal ranges predict changes in reef fish community structure during 8 years of extreme temperature variation. Diversity and Distributions 24:1036–1046. doi:10.1111/ddi.12753

Devictor V, Julliard R, Couvet D, Jiguet F. 2008. Birds are tracking climate warming, but not fast enough. Proceedings of the Royal Society B: Biological Sciences 275:2743–2748. doi:10.1098/rspb.2008.0878

Devictor V, van Swaay C, Brereton T, Brotons L, Chamberlain D, Heliölä J, Herrando S, Julliard R, Kuussaari M, Lindström Å, Reif J, Roy DB, Schweiger O, Settele J, Stefanescu C, Van Strien A, Van Turnhout C, Vermouzek Z, WallisDeVries M, Wynhoff I, Jiguet F. 2012. Differences in the climatic debts of birds and butterflies at a continental scale. Nature Clim Change 2:121–124. doi:10.1038/nclimate1347

Diamond SE, Frame AM, Martin RA, Buckley LB. 2011. Species’ traits predict phenological responses to climate change in butterflies. Ecology 92:1005–1012. doi:10.1890/10-1594.1

Duffy K, Gouhier TC, Ganguly AR. 2022. Climate-mediated shifts in temperature fluctuations promote extinction risk. Nat Clim Chang 12:1037–1044. doi:10.1038/s41558-022-01490-7

Engelhardt EK, Biber MF, Dolek M, Fartmann T, Hochkirch A, Leidinger J, Löffler F, Pinkert S, Poniatowski D, Voith J, Winterholler M, Zeuss D, Bowler DE, Hof C. 2022. Consistent signals of a warming climate in occupancy changes of three insect taxa over 40 years in central Europe. Global Change Biology 28:3998–4012. doi:10.1111/gcb.16200

ESRI. 2019. ArcGIS (Version 10.7.1).

Estrada A, Morales-Castilla I, Caplat P, Early R. 2016. Usefulness of Species Traits in Predicting Range Shifts. Trends in Ecology & Evolution 31:190–203. doi:10.1016/j.tree.2015.12.014

Franke S, Pinkert S, Brandl R, Thorn S. 2022. Modeling the extinction risk of European butterflies and odonates. Ecology and Evolution 12:e9465. doi:10.1002/ece3.9465

Franks SE, Pearce-Higgins JW, Atkinson S, Bell JR, Botham MS, Brereton TM, Harrington R, Leech DI. 2018. The sensitivity of breeding songbirds to changes in seasonal timing is linked to population change but cannot be directly attributed to the effects of trophic asynchrony on productivity. Global Change Biology 24:957–971. doi:10.1111/gcb.13960

Gaiji S, Chavan V, Ariño AH, Otegui J, Hobern D, Sood R, Robles E. 2013. Content assessment of the primary biodiversity data published through GBIF network: Status, challenges and potentials. Biodiversity Informatics 8. doi:10.17161/bi.v8i2.4124

García-Roselló E, Guisande C, Manjarrés-Hernández A, González-Dacosta J, Heine J, Pelayo-Villamil P, González-Vilas L, Vari RP, Vaamonde A, Granado-Lorencio C, Lobo JM. 2015. Can we derive macroecological patterns from primary Global Biodiversity Information Facility data? Global Ecology and Biogeography 24:335–347. doi:10.1111/geb.12260

Goulson D. 2019. The insect apocalypse, and why it matters. Current Biology 29:R967–R971. doi:10.1016/j.cub.2019.06.069

Grant PR, Grant BR, Huey RB, Johnson MTJ, Knoll AH, Schmitt J. 2017. Evolution caused by extreme events. Philosophical Transactions of the Royal Society B: Biological Sciences 372:20160146. doi:10.1098/rstb.2016.0146

Grewe Y, Hof C, Dehling DM, Brandl R, Brändle M. 2013. Recent range shifts of European dragonflies provide support for an inverse relationship between habitat predictability and dispersal. Global Ecology and Biogeography 22:403–409. doi:10.1111/geb.12004

Gutiérrez D, Wilson RJ. 2021. Intra- and interspecific variation in the responses of insect phenology to climate. Journal of Animal Ecology 90:248–259. doi:10.1111/1365-2656.13348

Hadfield JD. 2010. MCMC Methods for Multi-Response Generalized Linear Mixed Models: The MCMCglmm R Package. Journal of Statistical Software 33:1–22. doi:10.18637/jss.v033.i02

Hällfors MH, Heikkinen RK, Kuussaari M, Lehikoinen A, Luoto M, Pöyry J, Virkkala R, Saastamoinen M, Kujala H. 2024. Recent range shifts of moths, butterflies, and birds are driven by the breadth of their climatic niche. Evolution Letters 8:89–100. doi:10.1093/evlett/qrad004

Hällfors MH, Pöyry J, Heliölä J, Kohonen I, Kuussaari M, Leinonen R, Schmucki R, Sihvonen P, Saastamoinen M. 2021. Combining range and phenology shifts offers a winning strategy for boreal Lepidoptera. Ecology Letters 24:1619–1632. doi:10.1111/ele.13774

Harris RMB, Beaumont LJ, Vance TR, Tozer CR, Remenyi TA, Perkins-Kirkpatrick SE, Mitchell PJ, Nicotra AB, McGregor S, Andrew NR, Letnic M, Kearney MR, Wernberg T, Hutley LB, Chambers LE, Fletcher M-S, Keatley MR, Woodward CA, Williamson G, Duke NC, Bowman DMJS. 2018. Biological responses to the press and pulse of climate trends and extreme events. Nature Clim Change 8:579–587. doi:10.1038/s41558-018-0187-9

Hassall C. 2015. Odonata as candidate macroecological barometers for global climate change. Freshwater Science 34:1040–1049. doi:10.1086/682210

Hassall C, Thompson DJ. 2008. The effects of environmental warming on Odonata: a review. International Journal of Odonatology 11:131–153. doi:10.1080/13887890.2008.9748319

Iler AM, CaraDonna PJ, Forrest JRK, Post E. 2021. Demographic Consequences of Phenological Shifts in Response to Climate Change. Annual Review of Ecology, Evolution, and Systematics 52:221–245. doi:10.1146/annurev-ecolsys-011921-032939

IPCC. 2021. Climate Change 2021: The Physical Science Basis. Contribution of Working Group I to the Sixth Assessment Report of the Intergovernmental Panel on Climate Change. Cambridge University Press.

Jones CD, Kingsley A, Burke P, Holder M. 2008. Field guide to the dragonflies and damselflies of Algonquin Provincial Park and the surrounding area, 1st ed. The Friends of the Algonquin Park.

Kalkman VJ, Boudot J-P, Bernard R, De Knijf G, Suhling F, Termaat T. 2018. Diversity and conservation of European dragonflies and damselflies (Odonata). Hydrobiologia 811:269–282. doi:10.1007/s10750-017-3495-6

Kerr JT. 2020. Racing against change: understanding dispersal and persistence to improve species’ conservation prospects. Proceedings of the Royal Society B: Biological Sciences 287:20202061. doi:10.1098/rspb.2020.2061

Kerr JT, Pindar A, Galpern P, Packer L, Potts SG, Roberts SM, Rasmont P, Schweiger O, Colla SR, Richardson LL, Wagner DL, Gall LF, Sikes DS, Pantoja A. 2015. Climate change impacts on bumblebees converge across continents. Science 349:177–180. doi:10.1126/science.aaa7031

Kharouba HM, Algar AC, Kerr JT. 2009. Historically calibrated predictions of butterfly species’ range shift using global change as a pseudo-experiment. Ecology 90:2213–2222. doi:10.1890/08-1304.1

Kujala H, Vepsäläinen V, Zuckerberg B, Brommer JE. 2013. Range margin shifts of birds revisited – the role of spatiotemporally varying survey effort. Global Change Biology 19:420–430. doi:10.1111/gcb.12042

Lawlor JA, Comte L, Grenouillet G, Lenoir J, Baecher JA, Bandara RMWJ, Bertrand R, Chen I-C, Diamond SE, Lancaster LT, Moore N, Murienne J, Oliveira BF, Pecl GT, Pinsky ML, Rolland J, Rubenstein M, Scheffers BR, Thompson LM, van Amerom B, Villalobos F, Weiskopf SR, Sunday J. 2024. Mechanisms, detection and impacts of species redistributions under climate change. Nat Rev Earth Environ 5:351–368. doi:10.1038/s43017-024-00527-z

Lenoir J, Bertrand R, Comte L, Bourgeaud L, Hattab T, Murienne J, Grenouillet G. 2020. Species better track climate warming in the oceans than on land. Nat Ecol Evol 4:1044–1059. doi:10.1038/s41559-020-1198-2

Leroux SJ, Larrivée M, Boucher-Lalonde V, Hurford A, Zuloaga J, Kerr JT, Lutscher F. 2013. Mechanistic models for the spatial spread of species under climate change. Ecological Applications 23:815–828. doi:10.1890/12-1407.1

Littlefield CE, Krosby M, Michalak JL, Lawler JJ. 2019. Connectivity for species on the move: supporting climate-driven range shifts. Frontiers in Ecology and the Environment 17:270–278. doi:10.1002/fee.2043

Lustenhouwer N, Wilschut RA, Williams JL, van der Putten WH, Levine JM. 2018. Rapid evolution of phenology during range expansion with recent climate change. Global Change Biology 24:e534–e544. doi:10.1111/gcb.13947

Macgregor CJ, Thomas CD, Roy DB, Beaumont MA, Bell JR, Brereton T, Bridle JR, Dytham C, Fox R, Gotthard K, Hoffmann AA, Martin G, Middlebrook I, Nylin S, Platts PJ, Rasteiro R, Saccheri IJ, Villoutreix R, Wheat CW, Hill JK. 2019. Climate-induced phenology shifts linked to range expansions in species with multiple reproductive cycles per year. Nat Commun 10:4455. doi:10.1038/s41467-019-12479-w

MacLean SA, Beissinger SR. 2017. Species’ traits as predictors of range shifts under contemporary climate change: A review and meta-analysis. Global Change Biology 23:4094–4105. doi:10.1111/gcb.13736

Mair L, Hill JK, Fox R, Botham M, Brereton T, Thomas CD. 2014. Abundance changes and habitat availability drive species’ responses to climate change. Nature Clim Change 4:127–131. doi:10.1038/nclimate2086

McCauley SJ, Hammond JI, Frances DN, Mabry KE. 2015. Effects of experimental warming on survival, phenology, and morphology of an aquatic insect (Odonata). Ecological Entomology 40:211–220. doi:10.1111/een.12175

McCauley SJ, Hammond JI, Mabry KE. 2018. Simulated climate change increases larval mortality, alters phenology, and affects flight morphology of a dragonfly. Ecosphere 9:e02151. doi:10.1002/ecs2.2151

Menzel A, Sparks TH, Estrella N, Koch E, Aasa A, Ahas R, Alm-Kübler K, Bissolli P, Braslavská O, Briede A, Chmielewski FM, Crepinsek Z, Curnel Y, Dahl Å, Defila C, Donnelly A, Filella Y, Jatczak K, Måge F, Mestre A, Nordli Ø, Peñuelas J, Pirinen P, Remišová V, Scheifinger H, Striz M, Susnik A, Van Vliet AJH, Wielgolaski F-E, Zach S, Zust A. 2006. European phenological response to climate change matches the warming pattern. Global Change Biology 12:1969–1976. doi:10.1111/j.1365-2486.2006.01193.x

Nagelkerke NJD. 1991. A Note on a General Definition of the Coefficient of Determination. Biometrika 78:691–692. doi:10.2307/2337038

Neate-Clegg MHC, Tonelli BA, Tingley MW. 2024. Advances in breeding phenology outpace latitudinal and elevational shifts for North American birds tracking temperature. Nat Ecol Evol 8:2027–2036. doi:10.1038/s41559-024-02536-z

New M, Lister D, Hulme M, Makin I. 2002. A high-resolution data set of surface climate over global land areas. Climate Research 21:1–25. doi:10.3354/cr021001

Ni M, Vellend M. 2021. Space-for-time inferences about range-edge dynamics of tree species can be influenced by sampling biases. Global Change Biology 27:2102–2112. doi:10.1111/gcb.15524

Novella-Fernandez R, Brandl R, Pinkert S, Zeuss D, Hof C. 2023. Seasonal variation in dragonfly assemblage colouration suggests a link between thermal melanism and phenology. Nat Commun 14:8427. doi:10.1038/s41467-023-44106-0

Pacifici M, Rondinini C, Rhodes JR, Burbidge AA, Cristiano A, Watson JEM, Woinarski JCZ, Di Marco M. 2020. Global correlates of range contractions and expansions in terrestrial mammals. Nat Commun 11:2840. doi:10.1038/s41467-020-16684-w

Parmesan C, Ryrholm N, Stefanescu C, Hill JK, Thomas CD, Descimon H, Huntley B, Kaila L, Kullberg J, Tammaru T, Tennent WJ, Thomas JA, Warren M. 1999. Poleward shifts in geographical ranges of butterfly species associated with regional warming. Nature 399:579–583. doi:10.1038/21181

Parmesan C, Yohe G. 2003. A globally coherent fingerprint of climate change impacts across natural systems. Nature 421:37–42. doi:10.1038/nature01286

Paulson D. 2012. Dragonflies and Damselflies of the East. Princeton University Press.

Pearse WD, Davis CC, Inouye DW, Primack RB, Davies TJ. 2017. A statistical estimator for determining the limits of contemporary and historic phenology. Nat Ecol Evol 1:1876–1882. doi:10.1038/s41559-017-0350-0

Pinkert S, Clausnitzer V, Acquah-Lamptey D, De Marco P, Johansson F. 2022. Odonata as focal taxa for biological responses to climate change In: Cordoba-Aguilar A, Beatty C, Bried J, editors. Dragonflies and Damselflies: Model Organisms for Ecological and Evolutionary Research. Oxford University Press. p. 0. doi:10.1093/oso/9780192898623.003.0027

Pinsky ML, Worm B, Fogarty MJ, Sarmiento JL, Levin SA. 2013. Marine Taxa Track Local Climate Velocities. Science 341:1239–1242. doi:10.1126/science.1239352

Platts PJ, Mason SC, Palmer G, Hill JK, Oliver TH, Powney GD, Fox R, Thomas CD. 2019. Habitat availability explains variation in climate-driven range shifts across multiple taxonomic groups. Sci Rep 9:15039. doi:10.1038/s41598-019-51582-2

Powney GD, Brooks SJ, Barwell LJ, Bowles P, Fitt RNL, Pavitt A, Spriggs RA, Isaac NJB. 2014. Morphological and Geographical Traits of the British Odonata. Biodivers Data J e1041. doi:10.3897/BDJ.2.e1041

Powney GD, Cham SSA, Smallshire D, Isaac NJB. 2015. Trait correlates of distribution trends in the Odonata of Britain and Ireland. PeerJ 3:e1410. doi:10.7717/peerj.1410

Pyke GH, Ehrlich PR. 2010. Biological collections and ecological/environmental research: a review, some observations and a look to the future. Biological Reviews 85:247–266. doi:10.1111/j.1469-185X.2009.00098.x

R Core Team. 2019. R: A language and environment for statistical computing.

Rapacciuolo G, Ball-Damerow JE, Zeilinger AR, Resh VH. 2017. Detecting long-term occupancy changes in Californian odonates from natural history and citizen science records. Biodivers Conserv 26:2933–2949. doi:10.1007/s10531-017-1399-4

Revell LJ. 2012. phytools: an R package for phylogenetic comparative biology (and other things). Methods in Ecology and Evolution 3:217–223. doi:10.1111/j.2041-210X.2011.00169.x

Richardson DM, Hellmann JJ, McLachlan JS, Sax DF, Schwartz MW, Gonzalez P, Brennan EJ, Camacho A, Root TL, Sala OE, Schneider SH, Ashe DM, Clark JR, Early R, Etterson JR, Fielder ED, Gill JL, Minteer BA, Polasky S, Safford HD, Thompson AR, Vellend M, Ehrlich PR. 2009. Multidimensional Evaluation of Managed Relocation. Proceedings of the National Academy of Sciences of the United States of America 106:9721–9724.

Robbirt KM, Davy AJ, Hutchings MJ, Roberts DL. 2011. Validation of biological collections as a source of phenological data for use in climate change studies: a case study with the orchid Ophrys sphegodes. Journal of Ecology 99:235–241. doi:10.1111/j.1365-2745.2010.01727.x

Rocha-Ortega M, Khelifa R, Sandall EL, Deacon C, Sánchez-Rivero X, Pinkert S, Patten MA. 2022. Linking traits to extinction risk in Odonata In: Cordoba-Aguilar A, Beatty C, Bried J, editors. Dragonflies and Damselflies: Model Organisms for Ecological and Evolutionary Research. Oxford University Press. p. 0. doi:10.1093/oso/9780192898623.003.0024

Rocha-Ortega M, Rodríguez P, Bried J, Abbott J, Córdoba-Aguilar A. 2020. Why do bugs perish? Range size and local vulnerability traits as surrogates of Odonata extinction risk. Proceedings of the Royal Society B: Biological Sciences 287:20192645. doi:10.1098/rspb.2019.2645

Román-Palacios C, Wiens JJ. 2020. Recent responses to climate change reveal the drivers of species extinction and survival. Proceedings of the National Academy of Sciences 117:4211–4217. doi:10.1073/pnas.1913007117

Rundle SD, Bilton DT, Abbott JC, Foggo A. 2007. Range size in North American Enallagma damselflies correlates with wing size. Freshwater Biology 52:471–477. doi:10.1111/j.1365-2427.2006.01712.x

Saino N, Ambrosini R, Rubolini D, von Hardenberg J, Provenzale A, Hüppop K, Hüppop O, Lehikoinen A, Lehikoinen E, Rainio K, Romano M, Sokolov L. 2010. Climate warming, ecological mismatch at arrival and population decline in migratory birds. Proceedings of the Royal Society B: Biological Sciences 278:835–842. doi:10.1098/rspb.2010.1778

Sandall EL, Pinkert S, Jetz W. 2022. Country-level checklists and occurrences for the world’s Odonata (dragonflies and damselflies). Journal of Biogeography 49:1586–1598. doi:10.1111/jbi.14457

Schuetz JG, Mills KE, Allyn AJ, Stamieszkin K, Bris AL, Pershing AJ. 2019. Complex patterns of temperature sensitivity, not ecological traits, dictate diverse species responses to climate change. Ecography 42:111–124. doi:10.1111/ecog.03823

Šigutová H, Pyszko P, Bílková E, Dolný A. 2025. Highly Conserved Ecosystems Facing Climate Change: Rapid Shifts in Odonata Assemblages of Central European Bogs. Global Change Biology 31:e70183. doi:10.1111/gcb.70183

Socolar JB, Epanchin PN, Beissinger SR, Tingley MW. 2017. Phenological shifts conserve thermal niches in North American birds and reshape expectations for climate-driven range shifts. Proceedings of the National Academy of Sciences 114:12976–12981. doi:10.1073/pnas.1705897114

Soroye P, Newbold T, Kerr J. 2020. Climate change contributes to widespread declines among bumble bees across continents. Science 367:685–688. doi:10.1126/science.aax8591

Souza KS, Fortunato DS, Jardim L, Terribile LC, Lima-Ribeiro MS, Mariano CÁ, Pinto-Ledezma JN, Loyola R, Dobrovolski R, Rangel TF, Machado IF, Rocha T, Batista MG, Lorini ML, Vale MM, Navas CA, Maciel NM, Villalobos F, Olalla-Tarraga MÂ, Rodrigues JFM, Gouveia SF, Diniz-Filho JAF. 2023. Evolutionary rescue and geographic range shifts under climate change for global amphibians. Front Ecol Evol 11. doi:10.3389/fevo.2023.1038018

Srivastava DS, Coristine L, Angert AL, Bontrager M, Amundrud SL, Williams JL, Yeung ACY, de Zwaan DR, Thompson PL, Aitken SN, Sunday JM, O’Connor MI, Whitton J, Brown NEM, MacLeod CD, Parfrey LW, Bernhardt JR, Carrillo J, Harley CDG, Martone PT, Freeman BG, Tseng M, Donner SD. 2021. Wildcards in climate change biology. Ecological Monographs 91:e01471. doi:10.1002/ecm.1471

Stephens PA, Mason LR, Green RE, Gregory RD, Sauer JR, Alison J, Aunins A, Brotons L, Butchart SHM, Campedelli T, Chodkiewicz T, Chylarecki P, Crowe O, Elts J, Escandell V, Foppen RPB, Heldbjerg H, Herrando S, Husby M, Jiguet F, Lehikoinen A, Lindström Å, Noble DG, Paquet J-Y, Reif J, Sattler T, Szép T, Teufelbauer N, Trautmann S, van Strien AJ, van Turnhout CAM, Vorisek P, Willis SG. 2016. Consistent response of bird populations to climate change on two continents. Science 352:84–87. doi:10.1126/science.aac4858

Suhonen J, Ilvonen JJ, Korkeamäki E, Nokkala C, Salmela J. 2022. Using functional traits and phylogeny to understand local extinction risk in dragonflies and damselflies (Odonata). Ecology and Evolution 12:e8648. doi:10.1002/ece3.8648

Sunday JM, Pecl GT, Frusher S, Hobday AJ, Hill N, Holbrook NJ, Edgar GJ, Stuart-Smith R, Barrett N, Wernberg T, Watson RA, Smale DA, Fulton EA, Slawinski D, Feng M, Radford BT, Thompson PA, Bates AE. 2015. Species traits and climate velocity explain geographic range shifts in an ocean-warming hotspot. Ecology Letters 18:944–953. doi:10.1111/ele.12474

Swaegers J, Janssens SB, Ferreira S, Watts PC, Mergeay J, McPeek MA, Stoks R. 2014. Ecological and evolutionary drivers of range size in Coenagrion damselflies. j evol Biol 27:2386–2395. doi:10.1111/jeb.12481

Urban MC. 2015. Accelerating extinction risk from climate change. Science 348:571–573. doi:10.1126/science.aaa4984

Usui T, Lerner D, Eckert I, Angert AL, Garroway CJ, Hargreaves A, Lancaster LT, Lessard J-P, Riva F, Schmidt C, van der Burg K, Marshall KE. 2023. The evolution of plasticity at geographic range edges. Trends in Ecology & Evolution 38:831–842. doi:10.1016/j.tree.2023.04.004

van der Laken P. 2021. ppsr: Predictive Power Score.

Waller JT, Svensson EI. 2017. Body size evolution in an old insect order: No evidence for Cope’s Rule in spite of fitness benefits of large size. Evolution 71:2178–2193. doi:10.1111/evo.13302

Waller JT, Willink B, Tschol M, Svensson EI. 2019. The odonate phenotypic database, a new open data resource for comparative studies of an old insect order. Sci Data 6:316. doi:10.1038/s41597-019-0318-9

Zattara EE, Aizen MA. 2021. Worldwide occurrence records suggest a global decline in bee species richness. One Earth 4:114–123. doi:10.1016/j.oneear.2020.12.005

Zografou K, Swartz MT, Adamidis GC, Tilden VP, McKinney EN, Sewall BJ. 2021. Species traits affect phenological responses to climate change in a butterfly community. Sci Rep 11:3283. doi:10.1038/s41598-021-82723-1

